# m6A is required for resolving progenitor identity during planarian stem cell differentiation

**DOI:** 10.1101/2021.07.27.453979

**Authors:** Yael Dagan, Yarden Yesharim, Ashley R. Bonneau, Schraga Schwartz, Peter W. Reddien, Omri Wurtzel

**Affiliations:** School of Neurobiology, Biochemistry, and Biophysics, The George S. Wise Faculty of Life Sciences, Tel Aviv University, 69978 Tel Aviv, Israel; Whitehead Institute for Biomedical Research, Cambridge, MA 02142, USA; Department of Biology, Massachusetts Institute of Technology, Cambridge, MA 02139, USA; Howard Hughes Medical Institute, Chevy Chase, MD 20815, USA; Department of Molecular Genetics, Weizmann Institute of Science, 7610001 Rehovot, Israel

## Abstract

Regeneration requires accurate production of missing cell lineages. Cell production is driven by changes to gene expression, which is shaped by multiple layers of regulation. Here, we find that the ubiquitous mRNA base-modification, m6A, is required for proper cell fate choice and cellular maturation in planarian stem cells (neoblasts). We mapped m6A-enriched regions in 7,600 planarian genes, and found that perturbation of the m6A pathway resulted in progressive deterioration of tissues and death. Using single cell RNA sequencing of >20,000 cells following perturbation of the pathway, we discovered that m6A negatively regulates transcription of histone variants, and that inhibition of the pathway resulted in accumulation of undifferentiated cells throughout the animal in an abnormal transcriptional state. Analysis of >1000 planarian gene expression datasets revealed that the inhibition of the chromatin modifying complex NuRD had almost indistinguishable consequences, unraveling an unappreciated link between m6A and chromatin modifications. Our findings reveal that m6A is critical for planarian stem cell homeostasis and gene regulation in regeneration.

## Introduction

Tissue maintenance and regeneration depend on the production of the necessary cell types. At a tissue scale, cell signaling balances cell production and contributes to selection of cell fates. Yet, at the scale of an individual cell, it is the regulation of gene expression that determines cell identity. Mechanisms that regulate gene expression of stem cells and other progenitors (e.g., dedifferentiated cells) are therefore central in the study of regeneration and tissue maintenance.

Recently, biochemical modifications to RNA molecules have emerged as regulators of gene expression. N6-methylation of adenosine (m6A) is a widespread modification to mRNAs, which is conserved from yeast to mammals. m6A modulates major aspects of RNA metabolism, including RNA stability, localization, and splicing^1^. The mRNAs are decorated with methyl groups installed by a methyltransferase complex (MTC), composed of the methyltransferase METTL3 and accessory proteins, including METTL14, KIAA1429, WTAP, Hakai, RBM15/RBM15B, and ZC3H13^1^. The methylated sites are recognized by several reader proteins including YTH-domain containing proteins affecting gene expression^2^. Studies in multiple organisms have shown that m6A, and many components of the m6A pathway, are essential for diverse processes: meiosis in the yeast *Saccharomyces cerevisiae*^3^, oogenesis and sex determination in *Drosophila melanogaster*^4, 5^, early development in mammal^6^, B-cell development^7^, and hematopoiesis^8^. m6A is therefore a key regulator of dynamic processes involving cellular proliferation and differentiation, and thus it could play an important role in tissue regeneration.

Planarians are flatworms that can regrow and support any adult tissue using a heterogeneous population of stem cells known as neoblasts^9^. Following injury, neoblasts rapidly divide and grow a blastema that forms the missing tissues. During homeostasis, neoblasts support existing tissues by continuously replacing them. The plasticity of individual neoblasts requires stringent regulation of gene expression^9–11^. Consequently, neoblast gene expression determines lineage selection^12, 13^, cell cycle progression^14^, and cellular migration^15^. Transcriptional regulation has been studied at multiple resolutions in planarians: extracellular signaling and other systemic signals that regulate neoblast biology have been studied in depth over the past decade. Epigenetic regulators that modify the chromatin landscape and alter fate determination were discovered^16^. Transcription factor expression that is associated with differentiation to distinct planarian lineages has been described^13^. However, molecular and functional studies of mRNA methylation have not been reported in planarians, and their impact on tissue homeostasis and regeneration is unknown.

Here, we analyzed the functions of planarian genes encoding components of the m6A-modification pathway (m6A genes) and studied the consequence of their inhibition. We mapped m6A-enriched regions in 7,600 genes expressed in a diversity of cell types and conditions. We identified multiple roles for the m6A pathway in neoblast biology, including regulation of cell fate choice, cell cycle control, and production of polyadenylated histone variants. Inhibition of m6A genes resulted in coordinated overexpression of adjacent genes in repetitive genomic regions, which were transcriptionally silenced in control animals. Systematic re-analysis of >1,000 planarian gene expression datasets revealed striking molecular similarities following silencing of the nucleosome remodeling and deacetylase complex (NuRD) component *CHD4*, suggesting a functional association between m6A and chromatin regulation. Single-cell RNA sequencing (scRNAseq) and confocal imaging revealed an accumulation of a previously undescribed class of cells following inhibition of m6A genes and of NuRD. Our study reveals that m6A is ubiquitous in planarians, establishes assays for studying m6A on a genomic scale, and reveals critical functions of m6A genes that are fundamental to regeneration and homeostasis.

## Results

### Components of m6A pathway are broadly expressed in planarians

We identified planarian genes that encode putative components of the MTC by searching for genes in the planarian genome encoding proteins similar to known MTC components in human, yeast, and fruit fly (Methods). We detected sequences corresponding to all of the canonical MTC components (Fig 1A, Supplementary Fig 1A). We identified six putative genes encoding m6A readers in the planarian genome by searching the YTH-domain signature in the predicted planarian proteome (Fig 1A, Supplementary Fig 1B). Orthologs of these putative YTH-encoding genes were found in six additional planarian species (Methods). By comparison, *Drosophila melanogaster* has two YTH-containing genes and vertebrates have five genes with a YTH-domain^17^, suggesting that the YTH gene family is expanded in planarians.

**Figure 1.**
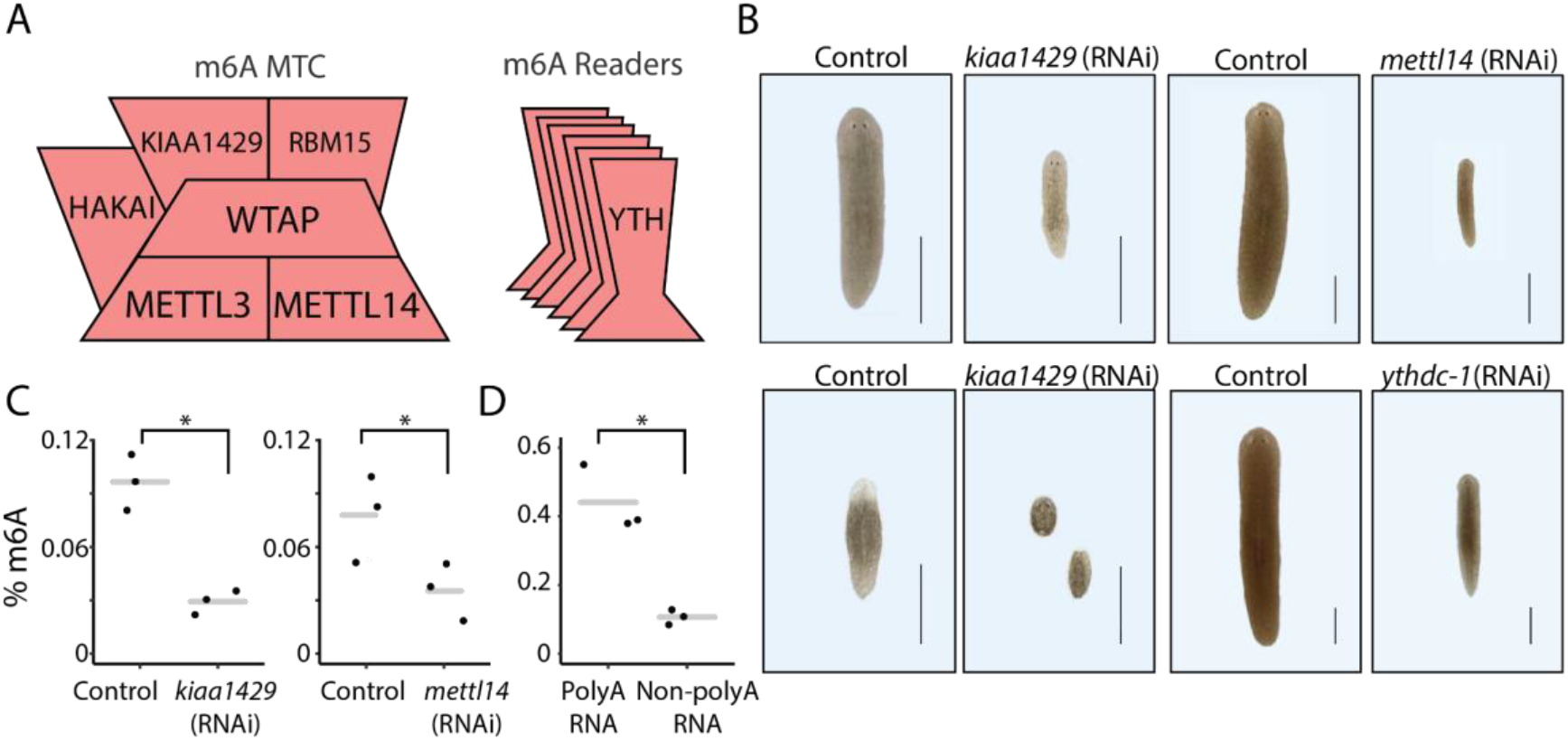
Expression of m6A genes is required for planarian regeneration and viability. (A) Components of the m6A MTC (left) and readers (right) that were identified in the planarian genome (Supplementary Fig 1A-B; Methods). (B) Effect of RNAi on planarian homeostasis and regeneration. Gene expression was inhibited by feeding the animals with dsRNA against the target gene (Methods). Phenotypes were observed in *kiaa1429* (RNAi) animals following five dsRNA feedings (top-left and bottom-left, homeostatic and 10 day regeneration phenotype, respectively) and following nine feedings of *mettl14* or *ythdc-1* dsRNA feedings (top and bottom, respectively). Experiments were performed at least three times with over 10 animals per experiment in each experimental group. Scale = 1 mm. (C-D) The relative abundance of m6A was estimated by using an anti-m6A-antibody (Supplementary Note 2), showing similar m6A levels compared to recent quantification of m6A in mammals^19;^ (C) Quantification of m6A on RNA isolated from *kiaa1429* (RNAi) and control animals following five RNAi feedings (left), or from *mettl14* (RNAi) and control animals following nine RNAi feedings (right), revealed a strong reduction in m6A levels following inhibition of gene expression (p < 0.05). (D) Comparison of m6A levels on polyadenylated and non-polyadenylated RNA has shown that polyadenylated RNA is highly enriched in m6A (p < 0.05).

We analyzed the expression of the genes that encode m6A-pathway components (m6A genes) across cell types in the planarian cell lineage atlas^18^. We found that m6A genes are expressed across multiple cell types, and particularly in neoblasts (Supplementary Fig 1C-D). The expression of genes that encode YTH-domain were expressed in largely non-overlapping cell types, suggesting that these putative readers might have distinct functions, which are activated in a cell type-specific manner (Supplementary Fig 1D). Only one YTH-domain gene, *ythdc-1*, was expressed broadly and across cells in multiple stages of differentiation. Analysis of MTC gene expression showed that MTC-encoding genes were expressed during the differentiation of multiple lineages and in multiple cell types (Supplementary Fig 1C). This suggests that m6A has a role in cellular differentiation in planarians, and that it might modulate different aspects of RNA metabolism using distinct m6A readers.

### m6A pathway activity is required for planarian regeneration and homeostasis

We studied the function of the m6A-pathway in planarians by systemically inhibiting the m6A genes using RNAi (Methods). Inhibition of components of the MTC encoded by *kiaa1429* and *mettl14* resulted in progressive reduction in body size, failure to uptake food, and eventually in lysis (Fig 1B, Supplementary Fig 1E-G). Inhibition of other MTC-encoding genes did not result in a penetrant phenotype, yet inefficient inhibition of gene expression could explain this result^20^ (e.g., of *mettl3*; Supplementary Fig 1H). Inhibition of the predicted m6A reader, *ythdc-1*, resulted in a similar phenotype (Fig 1B). Interestingly, *ythdc-1* is the only m6A reader gene that is broadly expressed across cell types and stages of lineage differentiation (Supplementary Fig 1D). Next, we tested the ability of animals to regenerate following the inhibition of the m6A pathway by feeding the animals with dsRNA and amputating them. Inhibition of both MTC encoding genes and *ythdc-1* eliminated the ability of the animals to regenerate. For example, amputated *kiaa1429* (RNAi) animals did not produce a blastema and consequently died (Fig 1B). We tested if this rapid effect was an outcome of depletion of m6A, or a nonspecific off-target effect of the RNAi. We tested the specificity of the *kiaa1429* (RNAi) phenotype by repeating the experiment using a non-overlapping dsRNA construct, and found comparable phenotypes (Supplementary Fig 1I-J; Supplementary Note 1). Next, we quantified the m6A levels of RNA from *kiaa1429* (RNAi), *mettl14* (RNAi), and control animals using an anti-m6A-antibody (Supplementary Note 2). We detected 3.3-fold and 2.2-fold reduction (p < 0.05) in the level of m6A (Fig 1C) following inhibition of *kiaa1429* and *mettl14*, respectively. Next, we tested if m6A was found on planarian mRNA, or on functional non-polyadenylated transcripts. Polyadenylated RNA was over four-fold enriched with m6A (Fig 1D; p < 0.05). These results demonstrate that (1) planarian m6A genes are required for regeneration and tissue homeostasis, (2) that the MTC genes are required for installing m6A, and (3) that m6A is enriched on mRNA.

### m6A methylation is abundant on planarian mRNAs

We next mapped m6A-enriched regions in planarian RNAs by using m6A-seq2^21^. m6A-seq2 uses an anti-m6A-antibody enrichment of methylated RNA followed by sequencing (Fig 2A; Supplementary Note 4). RNA was extracted from control animals, having normal m6A levels, and from *kiaa1429* (RNAi) animals, having depleted m6A (Fig 1C; Fig 2A). m6A-seq2 validated that RNA from *kiaa1429* (RNAi) animals was indeed m6A poor (Supplementary Fig 2A-D; Supplementary Note 4). We mapped the sequencing data to the planarian genome. Then, we used MeTPeak^22^ for detecting genomic regions with high read-mapping density (peaks) in the antibody-enriched RNA compared to input RNA (Fig 2A-B; Supplementary Note 5). m6A peaks represent regions likely enriched with m6A^23^. The read mapping was similar across all libraries (84-90%, SD=1.5%), without a significant difference between the mapping of m6A-enriched libraries and their input controls, and between *kiaa1429* (RNAi) or control RNA (Supplementary Note 5). Reads were aligned to >20k genes, producing a comprehensive map of the planarian m6A transcriptome (Supplementary Table 1). We detected 7,600 genes having m6A peaks enriched by >5-fold over input in the control m6A libraries (MeTPeak score < 1E-4). Many peaks were detectable following inhibition of *kiaa1429* but their enrichment score was significantly reduced (Fig 2B; Supplementary Fig 2B-C; Supplementary Table 1; p=2.5E-6). We divided the genes to bins based on their expression in the input libraries and examined m6A-enriched regions in each bin. The most highly expressed genes (bin 10) had fewer m6A peaks than genes in bins 6 to 9 (Fig 2C), an observation consistent with findings in mammalian systems^23, 24^. Interestingly, m6A was not detected on rRNA, in contrast with mammalian systems where two sites are installed via a mechanism independent of METTL3 (Supplementary Fig 2E). m6A peaks were strongly biased towards the 3’ end of transcripts, similar to the case in mammalian systems, a trend that was not observed in the input samples (Fig 2B). Interestingly, the peak length distribution in planarians was longer than observed in other organisms^25^ (Fig 2D). In most studied organisms m6A is installed on the conserved sequence motif DRACH (D=A/G/U; R=A/G; H=A/C/U)^26^. We searched for the DRACH motif in m6A-enriched sequences using CentriMo and performed de novo sequence motif discovery with STREME^27, 28^ on planarian m6A peak sequences. We could not detect significantly enriched sequence motifs. It is, in principle, possible that the lack of a detectable enrichment of the DRACH motif could be because of technical challenges in identifying enriched short motifs within the long peaks, or because of promiscuous enrichment of sequences by the antibody (Schwartz et al, Cell, 2013). However, we believe that this is unlikely to be the case, given the fact that the identified sites do exhibit many of the classical hallmarks of m6A: they are enriched towards the ends of genes, depleted from highly expressed genes, and most importantly, are highly depleted upon inhibition of *kiaa1429*. Our results thus suggest that the specificity of the m6A machinery in planarian systems may have diverged from the consensus sequence found in other Eukarya.

**Figure 2.**
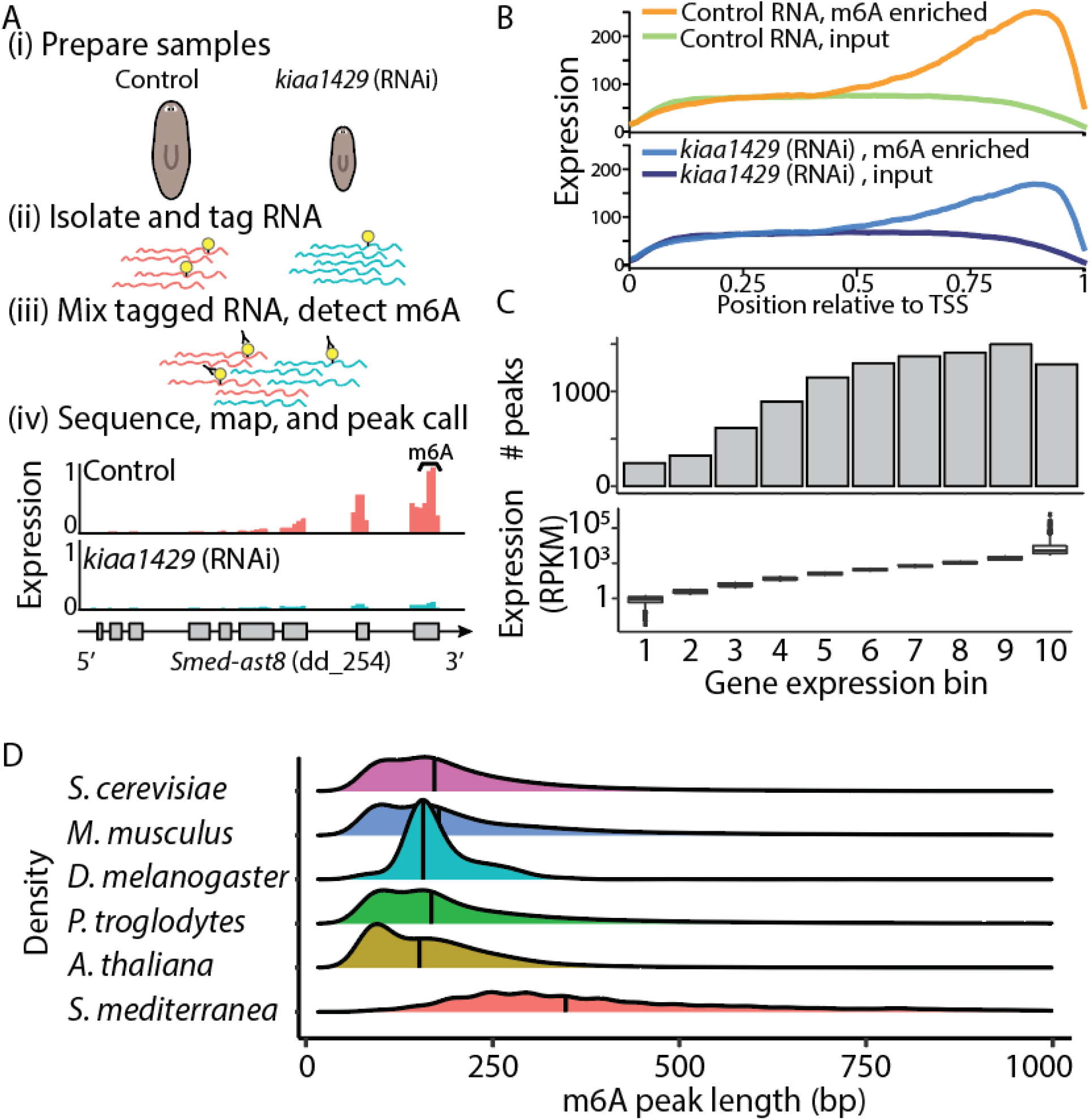
Transcriptome-wide mapping of m6A in planarians. (A) Outline of mapping of m6A on planarian RNA. RNA was extracted from control or *kiaa1429* (RNAi) animals in triplicates, with at least 10 animals in each replicate (i). Then, RNA was molecularly tagged with a barcoded RNA oligo (ii). Samples of barcoded RNA were mixed, and methylated RNA was enriched using an anti-m6A-antibody; input samples were sequenced without anti-m6A-antibody enrichment (iii). RNA was used for RNAseq library preparation, and resultant libraries were sequenced. Sequencing data was mapped to the planarian genome, and m6A-enriched regions (peaks) were detected using MeTPeak^22;^ coverage across *Smed-ast8* is shown as normalized RPKM that was scaled to 0-1 across samples (iv). m6A peaks are highly enriched at the 3’ end of the transcript. X-axis shows the relative position to the transcription start site (TSS); transcript lengths were normalized to 1000 bp. Y-axis is the average peak expression (normalized by RPKM) across 740 high-confidence peaks. Plot produced using the deepTools2 package^29^ (Supplementary Note 5). (C) The number of m6A peaks, which were determined using MetPeak^22,^ is shown (top) as a function of gene expression in each bin (bottom, box plot). (D) The distribution of the m6A peak length as detected by MeTPeak^22.^ Black line represents the average peak length. Processed data for organisms other than planarian were obtained from REPIC^25.^

### m6A genes regulate neoblast gene expression

The animal size reduction and lysis following inhibition of expression of m6A genes suggested that the animals were unable to produce cells normally. Failure to maintain and generate tissue is often a consequence of neoblast depletion or failure of neoblasts to differentiate^30–32^. We tested whether neoblasts were depleted following m6A-pathway inhibition by counting neoblasts by fluorescent in situ hybridization (FISH; Methods) using two canonical neoblast markers, *smedwi-1*^32^ and *h2b*^33^. However, the number of *smedwi-1*^+^ or *h2b*^+^ cells per unit area was similar following inhibition of *kiaa1429* gene expression (Fig 3A-B). Therefore, the phenotypes were not a consequence of neoblast depletion. Our FISH approach allowed counting neoblasts using neoblast markers, but not to measure gene expression levels. We used quantitative reverse transcription PCR (qPCR) for measuring *smedwi-1* and *h2b* expression level following RNAi on polyadenylated RNA (Fig 3C-D; Supplementary Note 6). We saw no significant change in *smedwi-1* gene expression in all of the tested RNAi conditions. By contrast, we found a significant upregulation of up-to 14 fold in *h2b* expression following inhibition of m6A genes. *h2b* was overexpressed even in conditions that did not produce a lysis phenotype (Fig 3D). The overexpression of *h2b* suggested that changes to neoblast gene expression preceded the development of the RNAi phenotype.

**Figure 3.**
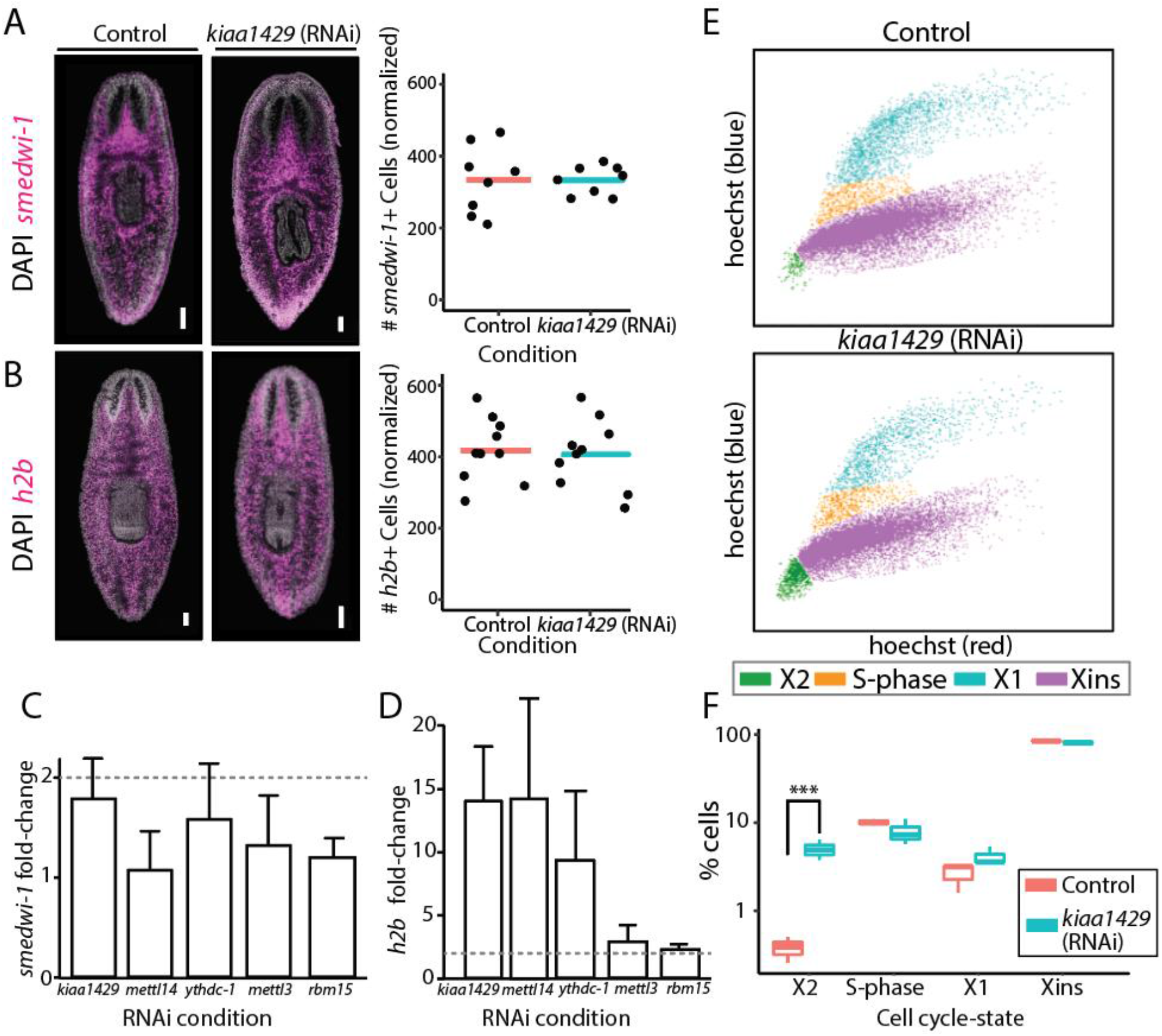
m6A pathway expression is not required for neoblast viability. (A-B) The number of neoblasts did not change significantly following inhibition of the MTC-gene *kiaa1429*. Shown is FISH of *smedwi-1* (A) and *h2b* (B) in control and *kiaa1429* (RNAi) animals (left). *smedwi-1*^+^ or *h2b*^+^ cells were counted following FISH (right; bar represents average normalized cell count; Methods). (C-D) Gene expression analysis of *smedwi-1* (C) and *h2b* (D). The expression of *smedwi-1* and *h2b* was determined in the different RNAi conditions compared to the expression of these genes in control animals (Supplementary Note 6). Error bars represent the 95% confidence interval, dashed line marks a two-fold change in expression. Measurements performed on biological triplicates. (E) Representative FACS plots of cells isolated from control (top) and *kiaa1429* (RNAi) animals (bottom) show an increase in cell abundance in the X2^36^ gate (green; Methods). (F) Quantification of the cells in each FACS gate in control and *kiaa1429* (RNAi) animals showed a six-fold increase in the abundance of cells in the X2 gate (*** - p < 0.001). FACS experiments were performed in triplicates. Scale = 100 µm.

Expression of *h2b* is a hallmark of neoblast cell-cycle progression^33–35^. We therefore tested whether the expression of m6A genes was required for normal cell cycle regulation. The relative abundance of cell cycle states was analyzed by fluorescence-activated cell sorting (FACS) of Hoechst-labeled planarian cells from control and *kiaa1429* (RNAi) animals (Fig 3E-F; Methods). G1 neoblasts and recently divided post-mitotic progenitors are found specifically in a FACS gate called X2^36^. Our FACS analysis (Fig 3E-F) revealed a six-fold increase in the number of cells in the X2 gate following *kiaa1429* RNAi, and more minor changes in the number of cells in the G2 gate (X1) or in fully differentiated cells (Xins). The increase in the X2 cell population could be a consequence of an increase (1) in G1 neoblasts; (2) in post-mitotic progenitors; or (3) in both. We considered these options. The FACS analysis revealed that there was a minor change in the abundance of S-phase and G2 cells (Fig 3E-F). In addition, FISH revealed that there was no absolute change in neoblast number following *kiaa1429* (RNAi), as determined by *smedwi-1*^+^ or *h2b*^+^ cell labeling (Fig 3A). Taken together, these data suggested that the population size of G1/S/G2 neoblasts did not change dramatically. Therefore, the increase in X2 cell abundance was most likely a consequence of an increase in the number of immature *smedwi-1*^-^ post-mitotic progenitors, and not of G1 neoblasts, which are *smedwi-1*^+^.

### m6A genes regulate cell-type specific gene expression programs

The increase in post-mitotic progenitors following the inhibition of m6A pathway components may have affected multiple cellular lineages and gene expression programs. We profiled the emergence of gene expression changes following inhibition of *mettl14* and *ythdc-1* after four, six, and eight RNAi feedings (Fig 4A; Supplementary Table 2; Supplementary Note 7). The phenotype of *kiaa1429* (RNAi) animals developed more rapidly, and therefore we measured gene expression changes only following five RNAi feedings (Fig 4B). The gene expression changes in the different RNAi conditions were remarkably similar with correlation > 0.7 in all time points (Fig 4A-C). The sequencing validated the specificity of our RNAi, and corroborated our qPCR results (Fig 4A-B, Supplementary Fig 3A-D). This indicated that the phenotypes were a consequence of m6A depletion and not m6A-independent functions of the MTC.

**Figure 4.**
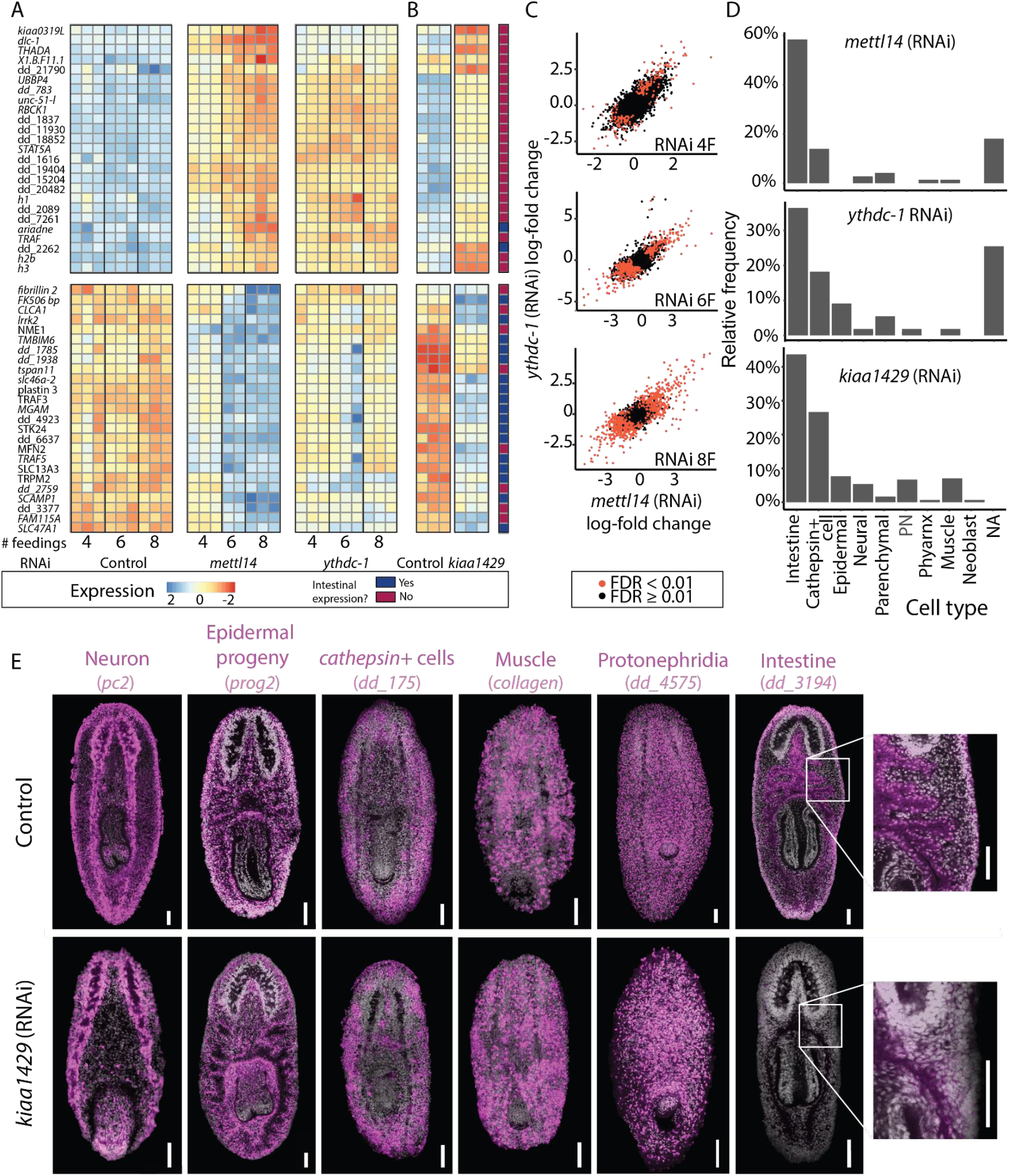
Gene expression changes following inhibition of m6A genes. (A-B) Heatmap of genes that changed their expression following inhibition of m6A genes. Shown are the expression levels of the top 25 overexpressed and 25 underexpressed genes (top and bottom, respectively; Supplementary Table 2) following six *mettl14* dsRNA feedings, at different conditions and time points. Rows and columns represent genes and samples, respectively. Blue and red, low to high gene expression (row-normalized z-score). Genes that were previously determined to be expressed in the intestine^39^ were highlighted in dark blue, or otherwise in dark red. (C) Correlation between gene expression changes between *mettl14* (RNAi) and *ythdc-1* (RNAi) compared to their controls. Each colored dot represents a gene (expression > 1 transcripts per million, TPM; red and black, significant and non-significant change in gene expression compared to controls, respectively). (D) The proportion of genes that were downregulated following RNAi, and were found to be expressed in a single cell type. Shown is an analysis based on the top 300 genes that were down-regulated following RNAi (six dsRNA feedings for *mettl14* or *ythdc-1*). (E) Detection of major planarian cell types following inhibition of *kiaa1429* by RNAi using previously established cell-type-specific gene expression markers^39.^ The distribution of most major planarian cell types was comparable between the control (top) and *kiaa1429* (RNAi) animals (bottom). A striking exception was the reduction in intestine cells (right), which severely affected the intestine branching morphology. Scale = 100 µm.

The presence of m6A has been previously associated with a high transcript turnover^37, 38^. Therefore, we predicted that inhibition of the m6A pathway would result in an overexpression of transcripts at early stages of the phenotype emergence. Indeed, following four RNAi feedings, most differentially expressed genes were upregulated: 77% and 83% for *mettl14* (RNAi) and *ythdc-1* (RNAi), respectively (Supplementary Fig 3E; Supplementary Table 2). By contrast, at late time points, most differentially expressed genes were downregulated (Supplementary Fig 3E), which likely reflected the indirect effects of m6A gene inhibition, such as depletion of a cell type (Supplementary Fig 3E).

We annotated differentially-expressed genes using the planarian cell-type transcriptome atlas^39^. The majority of the upregulated genes following RNAi were not cell type-specific (Supplementary Table 2), nor did they share a common function^40^, or had association with the injury response^41^. By contrast, a major fraction of the downregulated genes were cell type-specific and were normally expressed in the planarian intestine and intestine-associated neoblasts^18, 39^ (Fig 4A-B, D). For example, the transcription factor *nkx2.2*, which is required for specification of intestine cells^42^, was downregulated following RNAi, and similarly, *activin-1* expression was reduced. The reduction in intestine-specific gene expression was observed already following four RNAi feedings, but it was exacerbated at later time points (Fig 4A-B; Supplementary Table 2).

To test whether the reduction in intestinal gene expression represented downregulation of a gene expression program or a reduction in intestine cell number, we analyzed by FISH *kiaa1429* (RNAi) and control animals using probes that label the planarian intestine (Fig 4E). We found a reduction in intestine cell labeling, and severe defects in intestinal branching morphology (Fig 4E), which suggested that the RNAi caused depletion of intestine cells. Indeed, following inhibition of m6A genes, animals did not uptake food (Supplementary Fig 1F). To test whether there was a major depletion in other cell types, we performed FISH using five other cell type markers for muscle, neuron, epidermal lineage, *cathepsin*+ cells, and protonephridia (Fig 4E). The general defect in *kiaa1429* (RNAi) body morphology (Fig 1B) limited the ability to quantify the abundance of individual cell types. Yet, cell type-specific FISH labeling demonstrated that all major cell types were distributed across the animal, with the exception of intestine cells (Fig 4E). Importantly, cell types and lineages other than the intestine might have been affected by the RNAi, yet other factors, such as the cell type-specific turnover rate, might have caused the intestine to develop a significant defect more rapidly.

### m6A represses polyadenylated histone gene expression and silences repetitive DNA

The reduction in intestine cells together with the increase in the immature progenitor population that we observed by FACS in *kiaa1429* (RNAi) animals, suggested that there was a defect in cell production or maturation. We examined the identity of the overexpressed genes in our RNAseq data, which could potentially explain the overabundance of immature progenitors. Previous analysis of cycling planarian cells shows that histone gene expression changes dramatically during neoblast specialization^43^. We found that many histone encoding genes were overexpressed, indicating a broad change to neoblast cell state. For example, eight such genes were upregulated by over two fold in *mettl14* (RNAi) animals (Fig 5A; Supplementary Table 2). Particularly, *h2b* and *h3* were strongly overexpressed in all conditions (Fig 5A-B; Supplementary Fig 3C).

**Figure 5.**
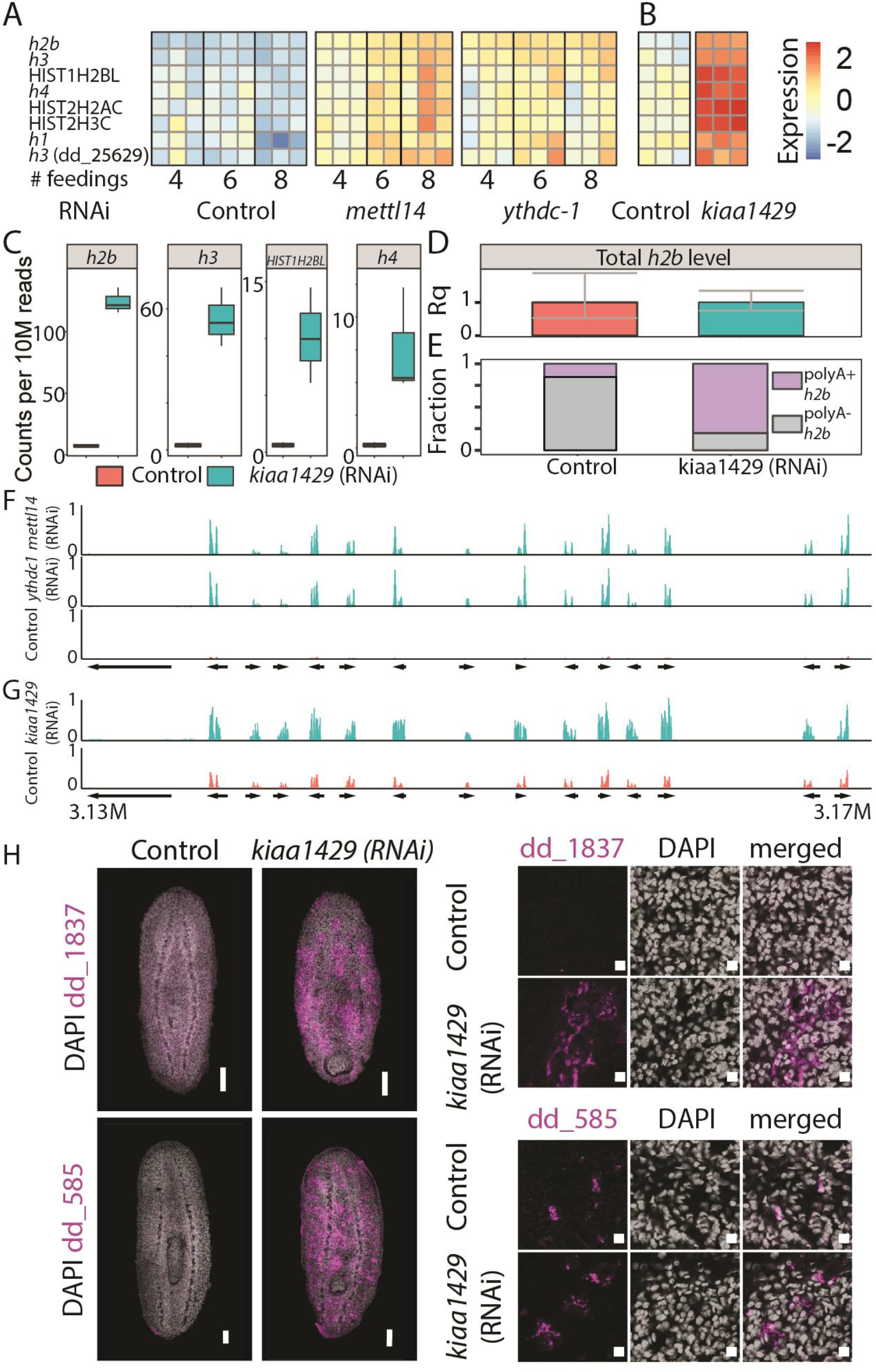
Overexpression of polyadenylated histone-encoding transcripts. (A-B) Heatmap showing eight histone-encoding genes that were upregulated following inhibition of m6A-pathway encoding genes. Rows and columns represent genes and samples, respectively. Blue and red, low to high gene expression (row-normalized z-score). Boxplots showing the normalized number of reads that map to a histone gene body and are paired with a read pair containing a polyA. Inhibition of *kiaa1429* resulted in a dramatic increase in the mapping to polyA in histone-encoding genes. (D) Quantification of the entire *h2b* transcript pool including polyA^+^ and polyA^-^ transcripts using qPCR (Methods; Supplementary Note 9). No significant change in the total *h2b* expression was observed following inhibition of *kiaa1429*. Error bars represent the 95% confidence interval. Experiment performed in biological triplicates. (E) Comparison of the polyA^+^ and polyA^-^ *h2b* transcripts showed an overabundance of polyA^+^ *h2b* in *kiaa1429* (RNAi) animals compared to controls. (F-G) Shown is a summary of gene mapping to a representative overexpressed gene neighborhood (contig: dd_Smes_g4_15). Arrows represent gene annotation, which was extracted from planmine^40.^ Panels show expression in TPM scaled to 0-1. (H) The overexpression of two representative genes from the upregulated genomic neighborhoods was validated by FISH. Cells expressing these genes were found throughout the animal in subepidermal and parenchymal layers (Left, Scale = 100 µm); higher magnification shows perinuclear expression (right, scale =10 µm).

Canonical histone-encoding genes are transcribed without a polyA tail^44^. Our RNAseq library preparation protocol selects polyadenylated RNA by using poly-dT beads (Methods). Therefore, non-polyadenylated RNA should have been vastly excluded from the RNAseq libraries. We hypothesized that the high prevalence of histone-encoding gene transcripts might have represented an aberrant expression of polyadenylated histone isoforms, which were undetectable when the m6A pathway functioned normally.

We tested this hypothesis by searching for polyadenylated histone transcripts in the RNAseq data. Our *kiaa1429* (RNAi) and control libraries were sequenced in a paired-end configuration (Supplementary Fig 4A). We reasoned that if *kiaa1429* inhibition activated polyadenylated histone transcription, we could detect RNAseq read pairs, where the first read in the pair mapped to a histone gene body, and the second read of the pair had a sequence that includes a polyT, which is the reverse-complement of the polyA tail (Supplementary Fig 4A; Supplementary Note 8). By contrast, in control samples, we expected to find fewer read pairs of this kind (Fig 5C; Supplementary Fig 4A-C). Indeed, polyadenylated histone transcripts were detectable almost exclusively in *kiaa1429* (RNAi) RNAseq compared to the control (Fig 5C). For example, we found a >17-fold increase in polyA read mapping to *h2b* following *kiaa1429* (RNAi).

The overexpression of polyadenylated histone RNAs following *kiaa1429* (RNAi) could represent an absolute increase in histone gene expression, or alternatively, a relative increase in polyadenylated histone transcripts compared to non-polyadenylated transcripts. We compared the total expression of *h2b* in *kiaa1429* (RNAi) and control animals using cDNA produced with random hexamers, which were used to reverse transcribe both polyadenylated and non-polyadenylated RNAs. We found that *h2b* expression levels were essentially identical (Fig 5D). This result indicated that the observed increase in *h2b* expression in the RNAseq data resulted from production of polyadenylated *h2b* transcripts and not from an absolute increase in *h2b* gene expression.

We estimated the proportion of polyadenylated *h2b* transcripts in RNA extracted from control or *kiaa1429* (RNAi) animals. We used cDNA produced with either random hexamers or a poly-dT primer, and validated that cDNA production using both methods was similarly efficient (Supplementary Fig 4D-E). In control animals, 15.4% of the *h2b* transcripts were polyadenylated. By comparison, following RNAi of *kiaa1429* 80.2% of the *h2b* transcripts were polyadenylated (Fig 5E, Supplementary Fig 4F-G). The shift towards production of polyA^+^ histone transcripts might reflect changes to cell cycle state, which is often associated with histone transcription^45, 46^.

The majority of overexpressed genes following inhibition of the m6A pathway were not associated with a distinct function (Fig 4A-B; Supplementary Table 2). However, some of the most overexpressed transcripts were annotated as pseudogenes (Supplementary Table 2). We mapped the RNAseq data to the planarian genome^40^, and inspected regions containing these overexpressed genes. Remarkably, overexpressed genes were highly enriched within genomic clusters (Fig 5F-G). These gene clusters were not expressed in control samples, but were dramatically induced following RNAi (Fig 5F-G; Supplementary Table 2). Importantly, the intergenic regions in these genomic neighborhoods were almost completely silenced, and therefore this overexpression was not a consequence of lack of transcriptional regulation over the entire genome segment (Fig 5F-G). We cloned two of these overexpressed genes, and performed FISH on *kiaa1429* (RNAi) and control animals (Fig 5H). FISH showed that these genes were indeed expressed only following *kiaa1429* RNAi, validating that expression of m6A genes was required for their silencing. We hypothesized that nuclear packaging might be altered. Analysis of nuclear labeling by DAPI of *kiaa1429* (RNAi) and control animals indicated that chromatin density was indeed different (Supplementary Fig 4H), with extraneous unlabeled nuclear regions, suggesting a change to nuclear packaging. In summary, inhibition of m6A genes resulted in broad changes to gene expression, including a coordinated overexpression of gene clusters that are normally silenced, together with an overabundance of polyadenylated histone transcripts, indicating an emergence of irregular cell states following inhibition of m6A genes.

### m6A pathway activity regulates cellular maturation

Our results indicated that m6A genes had different roles in different cell populations. We dissected the cell type-specific roles of m6A genes by using scRNAseq of >20,000 cells isolated from control and *kiaa1429* (RNAi) animals (Supplementary Note 10; Supplementary Fig 5A-D). Then, we used the Seurat framework^47^ and the planarian cell type atlas^18, 39^ for assigning cell type identities for all of the sequenced cells (Supplementary Table 3). We assigned the cells to 34 major groups, annotated the major planarian cell types, and analyzed gene expression changes following *kiaa1429* (RNAi) for each cell type (Fig 6A-B; Supplementary Table 3). We focused on several aspects of the m6A depletion phenotype: (1) Changes to neoblast gene expression and state; (2) Molecular characterization of the cells expressing the repetitive gene clusters; and (3) the effect of the RNAi on intestine cells.

**Figure 6.**
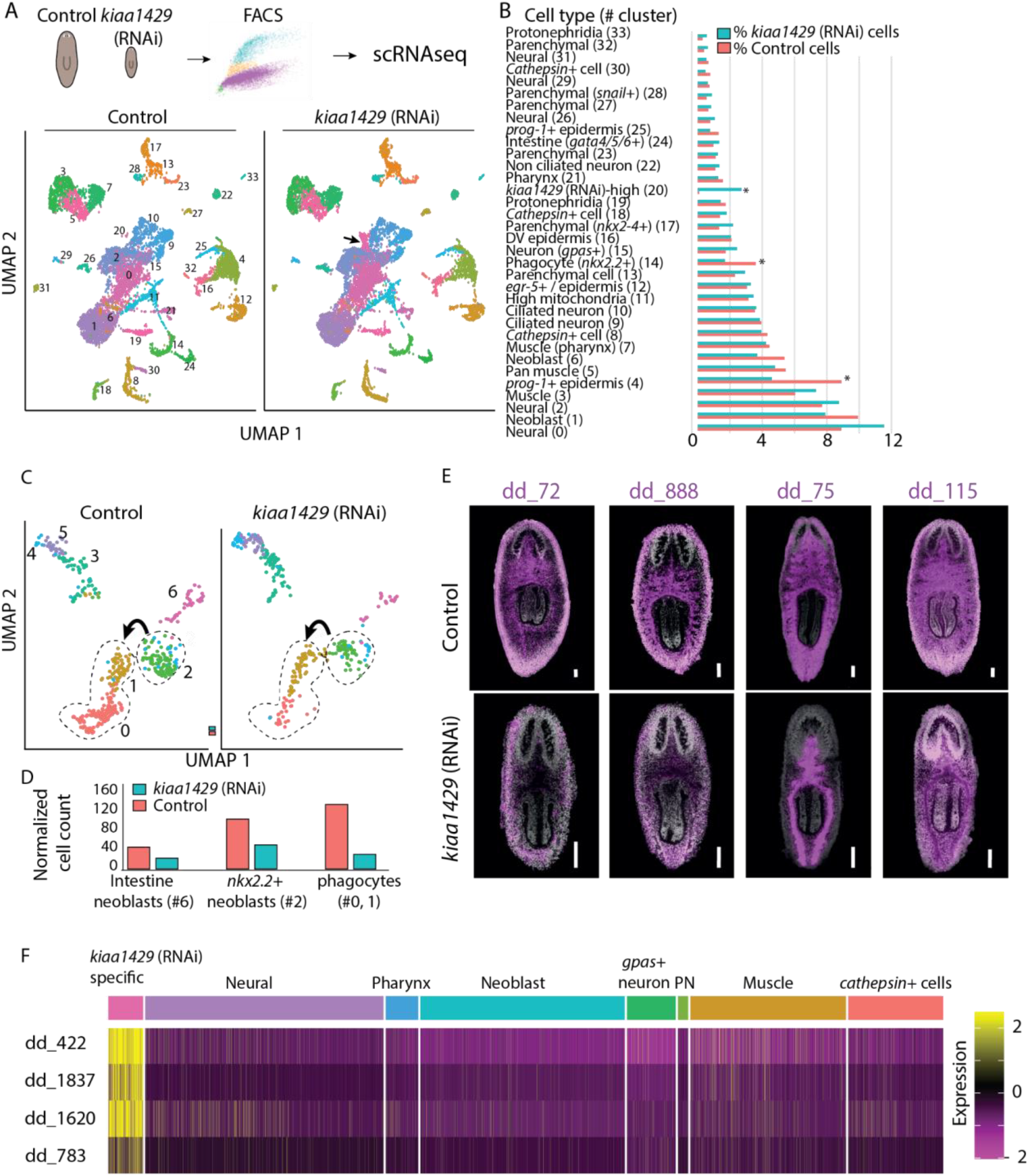
Cell type-specific regulation of gene expression by m6A. (A) Outline of scRNAseq experiment. Cells were isolated from dissociated animals, purified by FACS and subjected to scRNAseq (top). Analysis of cell populations detected in the scRNAseq and visualized by UMAP (dot represents cell, and color represents cluster association). Comparison of scRNAseq of *kiaa1429* (RNAi) and control reveals an expansion of a molecularly defined cell cluster (#20, black arrow). (B) Relative representation of clusters between *kiaa1429* (RNAi) and control animals. Most clusters were similarly represented between samples. However, depletion of phagocytic intestine cells, epidermal progenitors, and expansion of a *kiaa1429* (RNAi) cell cluster were detected. (C-D) Analysis of cells from the intestine lineage shows depletion in intestine neoblasts (gamma, #6), *nkx2.2*^+^ progenitors (#2), and differentiated intestine cells (#0, #1). Arrow indicates transition to differentiated cell state. Normalized cell count of intestine lineage cells are summarized (bottom). (E) FISH using markers for different intestine cell types^39^ demonstrated changes to gene expression and morphological defects following inhibition of *kiaa1429*. Scale = 100 µm. (F) scRNAseq heatmap produced using Seurat^47^ of four representative genes from the kiaa1429 (RNAi)-overexpressed genomic clusters. The major planarian cell types do not express these genes. By contrast, a *kiaa1429* (RNAi)-specific cell cluster is characterized by high expression of these genes. Rows and columns represent genes and cells, respectively. Purple to yellow, low to high gene expression. Top row shows assignment of cells to cell types based on gene expression (Supplementary Table 3) .

The number of *smedwi-1*+ cells was similar in control and *kiaa1429* (RNAi) animals. Expression of canonical neoblast genes, such as *smedwi-1*+, *Smed-vasa*, and *Smed-bruli*, was comparable, (Supplementary Fig 5E; Supplementary Table 3), indicating that the neoblast population indeed remained intact. By contrast, polyadenylated histone transcripts were broadly overexpressed in highly expressing *smedwi-1* cells following *kiaa1429* inhibition (Supplementary Fig 5F), suggesting a broad impact of the RNAi on the neoblast population transcriptional state. Neoblasts express lineage-specific genes during differentiation^48, 49^. We reclustered the neoblasts and compared lineage-specific gene expression in control and *kiaa1429* (RNAi) cells (Supplementary Fig 5G-H). We found two neoblast populations that were reduced by ∼40%: intestine-producing neoblast (*nkx2.2+*) and *PLOD*+ neoblasts (Supplementary Fig 5G-H), a class of neoblast that is transcriptionally associated with planarian muscle^39^. By contrast, there was an increase of 39%-63% in the number of neural-associated neoblasts, including in the *tspan-1*+ neoblasts^14, 50^ (Supplementary Fig 5G-H). These results suggested that m6A is required for lineage resolution, which if correct would impact the representation of differentiated cell populations. Examination of differentiated cells showed that several lineages were depleted in *kiaa1429* (RNAi) animals (Fig 6B; Supplementary Table 3). The strongest reduction in lineage representation (≥40%) was detected for intestinal phagocytes, consistent with the reduced representation of intestinal gene expression in bulk RNAseq (Fig 6C-D, 4A-B, Chi-squared p < 1E-15). Using FISH, we saw a reduction of multiple intestinal cell types (Fig 6E), and disruption to intestinal morphology^39, 51^. The intestine lineage was depleted at all stages of the lineage in the scRNAseq data (Fig 6C-D, Supplementary Fig 5I; Chi-squared p < 0.001; corrected with Bonferroni’s correction), starting at the intestine neoblasts (gamma), continuing with *nkx2-2*^+^/*smedwi-1*^+^ progenitors, and finally with the intestine phagocytes (Fig 6C-D, Supplementary Fig 5I). The dramatic decrease in differentiated cells (∼75%) following inhibition of *kiaa1429*, in comparison to a smaller decrease in intestine neoblasts (∼50%), might reflect an accumulated defect in cellular production. Indeed, the expression of intestine-specific-genes declined progressively in our RNAseq time courses (Fig 4A-B; Supplementary Table 2).

The increase in neural-associated neoblasts did not result in an overabundance of differentiated neural cells (Fig 6A-B; Supplementary Table 3). However, we identified a molecularly-defined population, which was overrepresented by >27-fold following *kiaa1429* inhibition (Fig 6B). This *kiaa1429*-specific cell cluster had characteristics of post-mitotic progenitors: (1) low-level, but detectable, expression of *smedwi-1* (Supplementary Fig 5J); (2) expression of genes associated with differentiation (Supplementary Fig 5K)^14^; and (3) expression of lineage-specific genes that are expressed early in differentiation (Supplementary Fig 5L; Supplementary Table 3). Remarkably, the cells in the cluster highly expressed genes that are associated with neurons and glia, such as *Smed-smad6/7-2* (Supplementary Fig 5K) and a protocadherin-like gene (dd_15376)^18, 39, 52^. These *kiaa1429* (RNAi)-specific cells were distinguished from essentially all other cells by expression of the repetitive gene clusters (Fig 6F, Supplementary Table 3). This result is consistent with the FISH analysis, which demonstrated a body-wide increase in cells expressing genes localized to the repetitive genomic clusters following inhibition of *kiaa1429* (Fig 5H). Taken together, these results support the hypothesis that inhibition of the m6A pathway leads to a decrease in differentiation of several cellular lineages, and the expansion of a molecularly-defined progenitor population.

### NuRD complex inhibition recapitulates the m6A inhibition phenotype

Multiple lines of evidence indicated that m6A gene expression was required for normal lineage-choice and differentiation. m6A is a broad regulator of cellular function, and the inhibition of the m6A pathway could produce this phenotype by affecting other processes directly or indirectly. To better understand the mechanistic basis for the m6A phenotypes, we searched for conditions that induce similar changes to gene expression. We mapped 1,080 published planarian RNAseq libraries to the planarian transcriptome^53^ and then compared gene expression changes in the published data with the RNAseq data that we generated following inhibition of the m6A genes (Supplementary Note 11; Supplementary Table 4).

Remarkably, this analysis revealed that inhibition of *Smed-CHD4*, which encodes a canonical component of the Nucleosome Remodeling and Deacetylase complex (NuRD), recapitulated all of the major aspects of the m6A phenotype (Supplementary Table 5): (1) overexpression of the repetitive gene clusters that were identified following RNAi of m6A genes (Fig 7A); (2) *CHD4* RNAi resulted in overexpression of polyadenylated histone-encoding transcripts (Fig 7B-D; Supplementary Fig 6A), as we observed in *kiaa1429* RNAi animals. (3) The expression of *smedwi-1* was unperturbed, indicating that the neoblast population size was unchanged (Supplementary Fig 6A); and (4) Strikingly, *CHD4* (RNAi) animals had reduced intestine gene expression, including transcription factors essential for intestinal cell differentiation, morphogenesis and function^42, 51^, as we previously observed following inhibition of the m6A genes (Fig 7E). We inhibited a different NuRD complex-encoding gene, *RbAp48*, which also resulted in overexpression of *h2b* (Supplementary Fig 6B). This further suggested that the molecular similarities detected following inhibition of m6A genes and *CHD4*, could be generalizable to other NuRD components, which is consistent with the roles of NuRD in transcriptional repression^54^. Interestingly, we analyzed previously published gene expression data from human and mouse of *CHD4* inhibition by shRNA or by conditional deletion and identified an overexpression of histone genes, such as *h2b*^55, 56^ (Supplementary Fig 6C-D; Supplementary Note 12). This indicated that regulation of histone gene expression by CHD4 is found in diverse animals.

**Figure 7.**
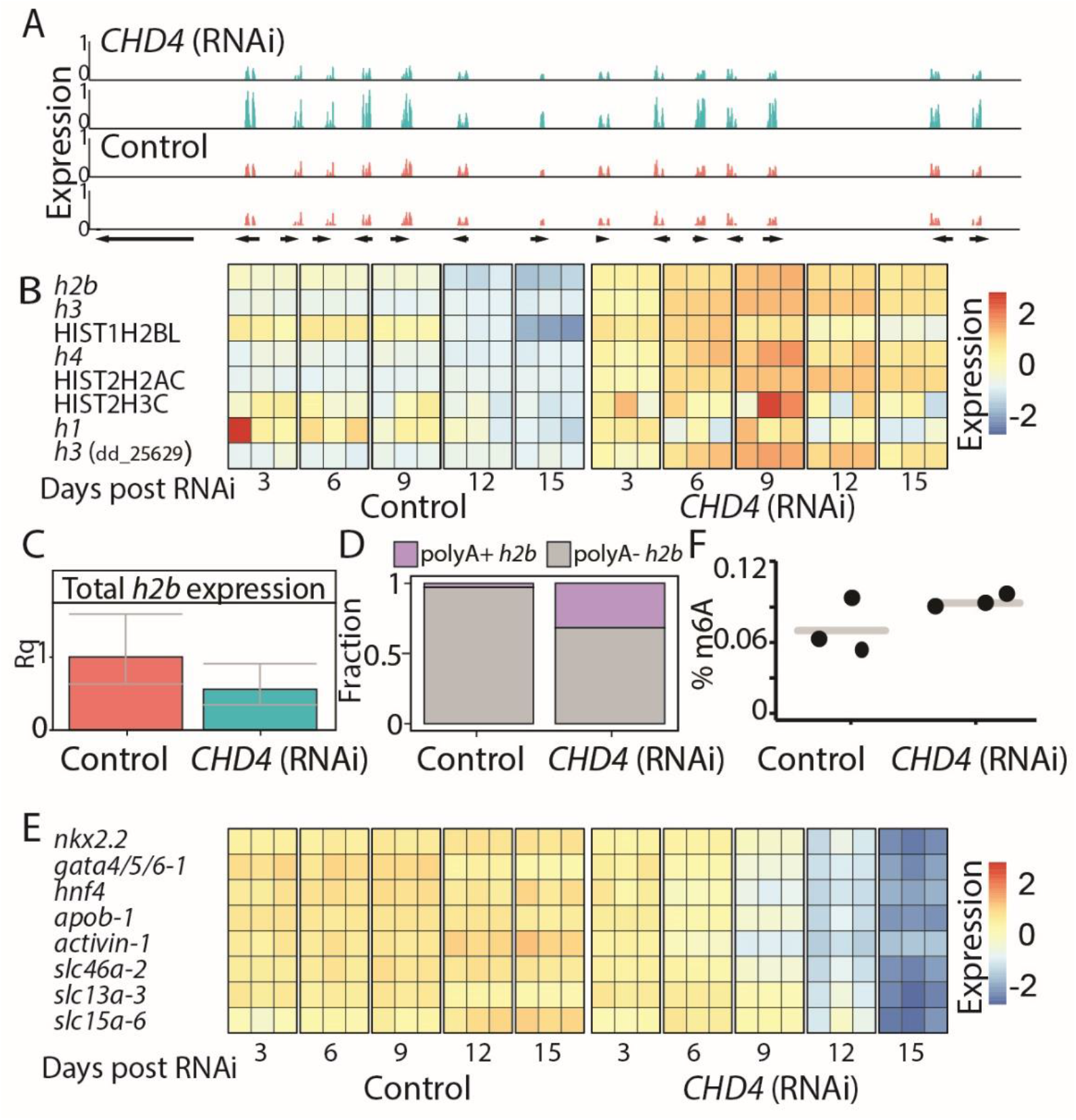
Inhibition of *CHD4* recapitulates the m6A-pathway inhibition. (A) Overexpressed gene neighborhood (contig: dd_Smes_g4_15:3.13-3.17 Mbp). Shown is the normalized and scaled gene expression in two *CHD4* (RNAi) samples, 12 days post feedings (top, blue) and in two corresponding control samples (bottom, red). (B) Heatmap showing the eight histone-encoding genes that were upregulated following inhibition of *CHD4*, at different time points, in published dataset^57.^ Rows and columns represent genes and samples, respectively. Blue and red, low to high gene expression (row normalized z-score). (C) Quantification of the entire *h2b* transcript pool including polyA^+^ and polyA^-^ transcripts using qPCR (Supplementary Note 9). No significant change in the absolute expression of *h2b* inhibition of *CHD4*. Error bars represent the 95% confidence interval. (D) Comparison of the polyA^+^ and polyA^-^ *h2b* expression level shows an increase in polyA^+^ *h2b* in *CHD4* (RNAi) animals compared to control animals. (E) Heatmap showing expression levels of intestine-specific TFs and genes following *CHD4* RNAi at different time points. Rows and columns represent genes and samples, respectively. Blue and red, low to high gene expression (row normalized z-score). (F) Quantification of m6A on RNA isolated from *CHD4* (RNAi) or control animals showed no significant change in abundance of m6A methylation.

We considered potential associations of m6A and NuRD, which might explain the similarities in the phenotypes (Fig 8): (1) NuRD and m6A regulate similar processes independently; (2) NuRD regulates m6A gene expression or m6A pathway activity; or (3) m6A regulates NuRD. We first examined whether NuRD activity was required for the expression of m6A genes, and found that the expression of m6A genes was unperturbed following inhibition of *CHD4*(Table S5). Then, we tested whether *CHD4* was required for m6A formation, and found that inhibition of *CHD4* did not deplete m6A (Fig 7F), indicating that NuRD is not necessary for m6A formation. Therefore, NuRD activity is unlikely to be required for the activity of the MTC. We next tested whether m6A genes regulated NuRD gene expression. Inhibition of m6A genes did not result in downregulation of *CHD4*, or any other NuRD-encoding components (Supplementary Table 2). This, of course, does not eliminate the possibility that the production or activity of one or more of NuRD components is regulated post-transcriptionally by m6A. Interestingly, *CHD4* (RNAi) animals developed phenotypes more rapidly than following inhibition of m6A genes. For example, lysis and changes to gene expression were observed following three RNAi feedings, in contrast to 8-10 RNAi feedings, which were required for lysis following *mettl14* (RNAi). This observation supports the possibility that m6A may progressively deteriorate NuRD activity, directly or indirectly, and indicates a potential link between m6A and chromatin regulation.

**Figure 8.**
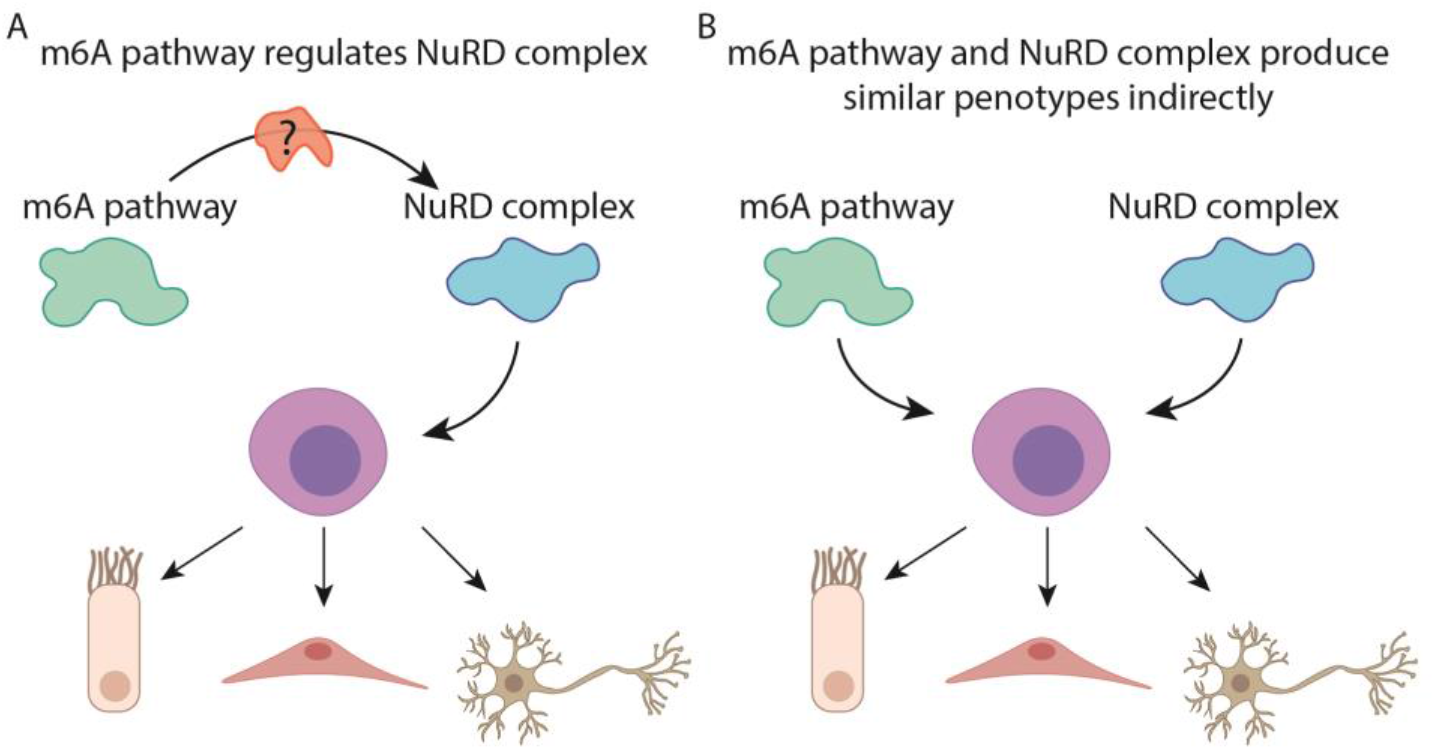
Potential models for interaction of m6A and NuRD. Shown are two models demonstrating the potential regulatory activity of m6A on neoblast differentiation. (A) m6A might regulate neoblast homeostasis by directly regulating NuRD or through a mediator that modulates NuRD function^64.^ (B) In the alternative model m6A and NuRD regulate similar processes, in parallel, without an association between the pathways.

## Discussion

m6A has been implicated in myriad regulatory roles in diverse biological systems^4, 8, 20, 58^. Given the essentiality of the MTC, research in mammalian systems has been limited to studying the role of this complex either in cell lines^6, 19, 20, 23^, or upon selective depletion of this complex in specific tissues or cell types, which has provided a partial and fragmented view on the roles played by this modification. Obtaining a global view on the functions and underlying mechanisms of action of m6A can hugely benefit from systemic, organism-wide perturbations of the MTC, as have been conducted in a highly limited number of model organisms to date, most of which in species evolutionary distant from human such as yeast and plants^3, 4, 24, 59, 60^. Our study adds to these efforts and introduces a systematic analysis of the MTC function in an adult Lophotrochozoan, planarian, which is a widely used model for studying regeneration.

Among the major questions for which we lacked a general understanding to date is the nature of the relationship between methylation writers and readers. All organisms surveyed to date have only a single ‘writer’ machinery, but organisms vary widely in terms of the number of readers that they have, ranging from one detected reader in yeast^3^ to 13 in plants^61^. Oftentimes, deletion of the writers is associated with a severe phenotype, whereas readers are associated with much milder counterparts. In some cases this has been attributed to redundancy between readers^37, 62, 63^, but in most cases it cannot be ruled out that the full function of the writers is not mediated by readers, but instead by other machineries^61^, potentially also methylation independent ones. In this regard, our findings mark a relatively rare exception – alongside with a previous report in drosophila^4^, and some recently reported findings in mouse embryonic stem cells (mESC) – in which the full phenotype associated with deletion of a writer is phenocopied via deletion of one reader. This strongly suggests a methylation-dependent function, and demonstrates that this function is likely to be mediated entirely via one reader. In parallel, these findings raise questions regarding the role played by the additional readers encoded in the planarian genome, whose disruption fails to give rise to a discernible phenotype. The absence of a phenotype may either be technical, or due to functional redundancy, and will be assessed in future studies.

We profiled the distribution of m6A in the planarian transcriptome, and identified 7,600 enriched regions in a diversity of cell types. The m6A-enriched regions were found mostly near the 3’ end of the transcripts, as previously reported in other systems^23, 24^. This suggested that the role of m6A in planarians is similar to other systems. However, we identified several unique features of planarian m6A-enriched regions: (1) they are longer, on average, than reported m6A-enriched regions in other organisms^25^; and (2) we could not detect the conserved sequence motif in the m6A-enriched region. While in most studied organisms the m6A-specific motif is conserved^25^, there is some evidence that m6A might be installed on other sequences, specifically in plants and parasites^59, 60, 65–68^, although the dependency on the MTC and METTL3 might need to be further established. We considered two potential explanations for the lack of the sequence motif. First, detection of an enriched short motif in the long m6A-enriched regions is challenging, and therefore, the current data could be insufficiently powered. Alternatively, the long m6A-enriched regions might include one or more methylated adenosines, which are formed independent of a sequence motif. Single-nucleotide resolution based approaches for detection of m6A, such as miCLIP^69^, could further elucidate these findings.

Importantly, we found essentially identical phenotypes following inhibition of the MTC-encoding genes *mettl14*, *kiaa1429*, and of the putative m6A-reader, *ythdc-1*. The phenotypes were reminiscent of a stem cell deficiency in a regenerative adult organism^10^: decline in animal size, failure to regenerate, and eventually death. In a developmental system, similar defects would be detrimental. Indeed, elimination of m6A in developmental contexts has revealed that m6A is critical for early development, cell maturation, and differentiation^6, 20, 62^.

Integration of our analysis of the m6A phenotypes showed that m6A is required for production of the planarian intestine with more minor impact on other lineages. Defects in differentiation to certain lineages, but not necessarily to all, have been reported in embryonic stem cells (ESC)^20^. However, in addition to the defect in differentiation, the lack of m6A in planarians resulted in an emergence of a molecularly-defined cell type throughout the planarian body. These cells were undetectable in control animals, and they lacked characteristics of neoblasts: they expressed low levels of neoblast markers and had different morphology. Moreover, these abnormal cells expressed several genes that are broadly expressed in neural progenitors, such as a protocadherin-like gene (dd_15376) and *Smed-SMAD6/7-2*. The scRNAseq clustering analysis positioned these cells adjacent to other neural cells. Therefore, m6A, in planarians, is required for resolving the identity of the differentiating progenitors.

Recently, an analysis of mESCs has found that deletion of *Mettl3* promotes an open state of the chromatin by regulating a class of chromatin associated RNAs^70^. In addition, *Mettl3* regulates self-renewal of neural stem cells by destabilizing transcripts that encode histone modifiers^71^. Analysis of planarian nuclei following inhibition of *kiaa1429* suggested that chromatin packaging was altered, which similarly to the findings in the mammalian system, may have profound effect on chromatin accessibility, and could be tested in future experiments. Moreover, we found that some of the most highly overexpressed genes, following inhibition of m6A genes, were localized to distinct genomic regions characterized by repetitive DNA. These regions were not expressed in control animals. Incorporation of the planarian cell type gene expression atlas^39^ with our m6A mapping data showed that 248 neoblast-associated transcripts are highly methylated, including many transcriptional regulators (Supplementary Table 1). Most importantly, inhibition of *CHD4*, a component of the NuRD complex, has revealed striking molecular similarities, including reduction in intestine cells and upregulation of the repetitive genomic regions. Potentially, m6A could affect the activity of chromatin regulators, or alternatively, regulate similar processes indirectly (Fig 8).

Interestingly, we did not detect NuRD complex genes that were highly methylated by m6A. Furthermore, inhibition of the m6A pathway did not affect NuRD complex gene expression. Similarly, inhibition of *CHD4* did not affect the levels of m6A on planarian RNAs, and did not result in a change to gene expression of canonical MTC components. Therefore, the striking similarities between the roles of m6A and NuRD, in planarians, could not be ascribed to simple transcriptional regulation. We therefore suggest two different models for the regulatory association of m6A and NuRD (Fig 8). In the first, m6A is required for proper activity of NuRD, potentially through a mediating molecule that is dependent on m6A methylation. Alternatively, m6A and NuRD regulate similar processes (e.g., chromatin accessibility) independently. Further study of neoblast-specific RNAs that are modified by m6A might reveal the identity of a mediating molecule, or distinguish between the two models.

In summary, we have mapped the distribution of m6A in the planarian transcriptome, characterized the roles of RNA-level modifications in planarian tissue maintenance and regeneration, and demonstrated that m6A is indispensable for regulation of neoblast homeostasis. Our study lays the foundation to a mechanistic analysis of planarian gene expression regulation and consequently a system-level understanding of planarian stem cell homeostasis.

## Methods

### Identification of putative m6A-pathway encoding genes

Sequences of genes encoding the MTC complex were extracted from the human, yeast, and fly genomes. Reciprocal BLAST search was performed with tblastx^72^ with e-value < 1x10^-5^ using the planarian transcriptome assembly^73^. Putative reader-encoding genes were identified by searching YTH domains in silico using hmmscan V3 with parameters [--noali --cut_nc --acc -- notextw]^74^. The resultant sequences were then searched in other planarian species in planmine^40^.

### Synthesis of Double-stranded RNA for RNAi experiments

Double stranded RNA (dsRNA) was synthesized as previously described^75^. Briefly, in vitro transcription (IVT) templates were prepared by PCR amplification of cloned target sequences using primers with 5’ flanking T7 promoter sequences. dsRNA was synthesized using the TranscriptAid T7 High Yield Transcription Kit (CAT K0441; Thermo Scientific). Reactions were incubated overnight at 37°C and then supplemented with RNase-free DNase for 30 minutes. RNA was purified by ethanol precipitation, and finally resuspended in 25 μl of ddH2O. RNA was analyzed on 1% agarose gel, and quantified by Qubit 4 (Thermo scientific) for validating a concentration higher than 5 μg/μl.

### cDNA synthesis from total RNA or polyadenylated RNA for qPCR

RNA samples were converted to cDNA and amplified using RevertAid H Minus First Strand cDNA Synthesis Kit (Thermo scientific; K1631). Random hexamers or oligo(dT) primers were used for obtaining cDNA from total RNA or from polyadenylated RNA, respectively. Resultant cDNA was used for qPCR analysis.

### Fluorescence-activated cell sorting (FACS)

Planarians were cut into small fragments using a blade. Tissue fragments were collected into calcium-free, magnesium-free medium plus 0.1% BSA (CMFB) and dissociated with collagenase l (1 mg/mL) by pipetting for 5-10 minutes. To reduce clumping at downstream steps, dissociated cells were strained through a 40 µm filter. Cells were centrifuged at 1250 rpm for 5 minutes at 4°C and resuspended in CMFB containing Hoechst 33342 (40 µL/mL) for 45 minutes at RT in the dark. Before loading into FACSAria II flow cytometer (Becton Dickinson), cells were labeled with propidium iodide (5 μg/ml) for cell viability detection. FACS gating was performed as previously described for planarian cell populations^36^.

### Preparation of RNA sequencing libraries

RNAseq libraries for *mettl14* (RNAi), *ythdc-1* (RNAi) and their control samples were prepared using Kapa stranded mRNAseq (KK8420) according to the manufacturer’s protocol.

### Planarian fixation for whole-mount assays

Fixation was performed as previously described^76^. Animals were killed with 5% N-acetyl-cysteine in PBS for 5 minutes, then fixed with 4% formaldehyde in PBSTx 0.1 for 20 minutes. Animals were then washed in PBSTx, 50:50 PBSTx:methanol and stored in methanol at -20°C until further analysis.

### Whole-mount fluorescence in situ hybridization

Fluorescence in situ hybridizations (FISH) was performed as previously described^76^ with minor modifications. Briefly, fixed animals were bleached, and treated with proteinase K (2 μg/ml) in 1x PBSTx. Following overnight hybridizations, samples were washed twice in pre-hyb solution, 1:1 pre-hyb-2X SSC, 2X SSC, 0.2X SSC, PBSTx. Subsequently, blocking was performed in 0.5% Roche Western Blocking Reagent and 5% inactivated horse serum in 1xPBSTx. Animals were incubated in an antibody overnight at 4°C (anti-DIG-POD, 1:1500). Post-antibody washes and tyramide development were performed as previously described^76^. Specimens were counterstained with DAPI overnight at 4°C (Sigma, 1 μg/ml in PBSTx).

### Cell counting following FISH in whole-mount planarian samples

FISH sample images were collected using a confocal microscope (Zeiss LSM800). Cell counting was performed on both controls and the experimental condition. Positive cells were counted using the “Cell Counter” component in ImageJ^77^, and the normalized positive cell count was calculated by measuring the area used for cell counting in each animal. Unless mentioned otherwise, the cells were counted in a rectangle starting at the posterior part of the brain and ending at the anterior of the pharynx in ventrally mounted animals.

### Data availability

High-throughput sequencing data produced in this project are available under accession PRJNA747686.

## Acknowledgements

We thank Sarit Edelheit for support with the m6A-seq2 protocol and Rami Khosravi for assistance with scRNAseq. We thank the Wurtzel lab for critical input. O.W. is supported by the Israel Science Foundation (grant 2039/18) and the European Research Council (no. 853640). O.W. is a Zuckerman Faculty Scholar. P.W.R. acknowledges NIH support (R01GM080639). P.W.R. is an Investigator of the Howard Hughes Medical Institute and an associate member of the Broad Institute of Harvard and MIT. S.S. is supported by the European Research Council (ERC) under the European Union’s Horizon 2020 research and innovation programme (grant agreement No. 714023). A.R.B. was supported by grant F32-GM128411 by the National Institute of General Medical Sciences.

## Author contributions

Y.D., Y.Y., A.R.B, P.W.R, and O.W. conceived and designed the experiments. Y.D., Y.Y., A.R.B and O.W. performed the experiments. Y.D., Y.Y., A.R.B, P.W.R, S.S, and O.W. analyzed the data. Y.D., Y.Y., and O.W. wrote and edited the paper with input from all authors.

