## Supplementary Material for "m6A is required for resolving progenitor identity during planarian stem cell differentiation"

### Table of contents

**Supplementary Note 1. Gene cloning and transformation** 4

Genes cloned in this study 4

**Supplementary Note 2. Measurement of m6A level in purified RNA** 6

**Supplementary Note 3. RNA extraction** 6

**Supplementary Note 4. mRNA m6A immunoprecipitation library preparation** 7

**Supplementary Note 5. Processing of m6A-seq2 library data** 8

**Inline m6a-seq2 barcode sequences** 8

**Mapping of library data to the planarian transcriptome** 9

**Supplementary Note 6. qPCR analysis of gene expression following RNAi** 10

**qPCR primers used in this study** 10

**Supplementary Note 7. Analysis of RNA sequencing gene expression changes** 11

**Supplementary Note 8. Detection of polyadenylated transcripts in RNAseq data** 12

**Supplementary Note 9. Quantification of polyadenylated h2b fraction by qPCR** 12

**Supplementary Note 10. Preparation of scRNAseq libraries following kiaa1429 (RNAi)** 12

**Supplementary Note 11. Systematic gene expression analysis of published planarian RNAseq data** 13

**Supplementary Note 12. Analysis of human and mouse gene expression data from CHD4 inhibition or conditional knockout** 13

**Supplementary Table 1. m6A-enriched regions across the planarian transcriptome** 14

**Supplementary Table 2. Gene expression changes following inhibition of m6A genes** 14

**Supplementary Table 3. scRNAseq of kiaa1429 (RNAi) and control animals** 14

**Supplementary Table 4. Published planarian high-throughput sequencing libraries compared to libraries produced here** 14

**Supplementary Table 5. Re-analysis of CHD4 (RNAi) RNAseq from Tu et al., 2015** 14

**Supplementary Figure 1** 15

**Supplementary Figure 2** 17

**Supplementary Figure 3** 19

**Supplementary Figure 4** 20

**Supplementary Figure 5** 22

**Supplementary Figure 6** 24

**Supplementary References** 25

### Supplementary Note 1. Gene cloning and transformation

Genes were amplified from planarian cDNA using gene-specific primers and cloned into pGEM-t vector using the manufacturer's protocol (Promega; CAT #A1360). Vectors were transformed into *E. coli* TOP10 (Thermo Fisher Scientific) by the heat-shock method. Briefly, 100 µl of bacteria was mixed with 5 µl of each of the cloned vectors, incubated on ice for 30 minutes, and then placed at 42°C for 45 seconds. Then, the transformed bacteria were supplemented with 350 µl of SOC medium, and following 1h of recovery at 37°C, the bacteria was plated on agarose plates containing 1:2000 Ampicillin, 1:200 Isopropylthio-b-D-galactoside (IPTG), and 1:625 5-bromo-4-chloro-3-indolyl-β- D-galactopyranoside (X-gal). Colonies were grown overnight at 37°C, and colonies were screened by colony PCR using M13F and M13R primers with the following PCR program: a. 5 minutes at 95°C; b. 34 cycles of 45 sec at 95°C, 60 sec at 55°C, and 2:30 minutes at 72°C; c. 10 minutes at 72°C; d. hold at 10°C. Reactions were analyzed by gel-electrophoresis, and correctly-sized gene products were grown overnight in Luria Broth (LB) medium, supplemented with 1:2000 Ampicillin at 37°C. Plasmids were purified from overnight cultures with the NucleoSpin Plasmid Miniprep Kit (Macherey-Nagel; CAT #740588). Cloned gene sequences were sequenced by Sanger sequencing.

#### Genes cloned in this study

| Contig | Comment | Sequence |
|---|---|---|
| dd_Smed_v4_4676_1 | Kiaa1429_F | CCGCTATCCGTTGTTATTATGCG |
| dd_Smed_v4_4676_1 | Kiaa1429_R | TCCGTAATCGTGGCCAGC |
| dd_Smed_v4_7282_1 | rbm15_F | GGGAGTTATCTGATGGTCAAAGA |
| dd_Smed_v4_7282_1 | rbm15_R | CCGGCACCGCTAACAGAA |
| dd_Smed_v4_6450_0_1 | mettl3_F | AATTGAAACCAGACGAAATGCA |
| dd_Smed_v4_6450_0_1 | mettl3_R | AGGTGTGTGGTTGCAGAGG |
| dd_Smed_v4_7426_1 | hakai_F | ACGGTCAATGTCTCGTAGCC |
| dd_Smed_v4_7426_1 | hakai_R | GCACGGCATCATGTTTAGG |
| dd_Smed_v6_2322_4 | wtap_F | CCCGATGAAATGGTGAAATC |

| | | |
|---|---|---|
| dd_Smed_v6_2322_4 | wtap_R | TCGGGATCATCCTCTTCATC |
| dd_Smed_v4_3491_0_1 | ythdc-1_F | CAGTCATCTCCCAATGTTGACG |
| dd_Smed_v4_3491_0_1 | ythdc-1_R | ACAAACCGCAATAATTGTAACCA |
| dd_Smed_v6_3194_0_1 | Intestine marker_F | GATCGAGGAAATCTTGACGAAC |
| dd_Smed_v6_3194_0_1 | Intestine marker_R | GTGAATACTTCAGGAGCCATCC |
| dd_Smed_v6_175_0_1 | *cathepsin*+ marker_F | ACGATCTCGGAAAACATTCG |
| dd_Smed_v6_175_0_1 | *cathepsin*+ marker_R | AGCCATCGTAGCTATTCCACA |
| dd_Smed_v6_701_0_1 | collagen F | GTGGAGAACTTGGAGCAAGC |
| dd_Smed_v6_701_0_1 | collagen R | AACACCAGCATCTCCTGGAC |
| dd_Smed_v6_4575_0_1 | Protonephridia_F | CAACCCCAACGCGATACAAT |
| dd_Smed_v6_4575_0_1 | Protonephridia_R | CAGAAAGAAAGAGCCTCCGC |
| dd_Smed_v4_4676_1 | *kiaa1429* non-overlapping_F | AGGACGTTTCCAGCAATGAG |
| dd_Smed_v4_4676_1 | *kiaa1429* non-overlapping_R | CATGGTTCGCCTTGGATTAG |
| dd_Smed_v6_2331_0_1 | *CHD4*_F | CGCGAGCATTTCATCTTGTA |
| dd_Smed_v6_2331_0_1 | *CHD4*_R | AAACGATGGGCTTCATCAAC |
| dd_Smed_v6_2065_0_1 | *rbAp48*_F | GAATGGCCAAGCTTAACTGC |
| dd_Smed_v6_2065_0_1 | *rbAp48*_R | TTTGGGCTCCAACTGAAATC |
| dd_Smed_v6_115_0_1 | Intestine marker_F | CAGTGCTTGCCGTCTGTCTA |
| dd_Smed_v6_115_0_1 | Intestine marker_R | TAGCAACCAGTGCATTGAGC |
| dd_Smed_v4_72_0_1 | Intestine marker_F | AATGTTGGGATGCTGCAGTT |
| dd_Smed_v4_72_0_1 | Intestine marker_R | CAAAACGCAGGGCTCATACT |
| dd_Smed_v4_75_0_1 | Intestine marker_F | TGCCGTTATGAACATGATTTTCG |
| dd_Smed_v4_75_0_1 | Intestine marker_R | ACACAAAATATCGCATCCTGCC |

| | | |
|---|---|---|
| dd_Smed_v4_888_0_1 | Intestine marker_F | TCTTGGACTTCATCGACTTTCT |
| dd_Smed_v4_888_0_1 | Intestine marker_R | AACTGGTTTTCGTTGTCTACAAA |
| dd_Smed_v6_1837_0_1 | *kiaa1429*-cluster #1 F | TTGGAACAGACCACTGGTGA |
| dd_Smed_v6_1837_0_1 | *kiaa1429*-cluster #1 R | AACGACGACCTTTCCAACTG |
| dd_Smed_v6_585_0_1 | *kiaa1429*-cluster #2 F | ACAAGTGCAATGCCCGTAAC |
| dd_Smed_v6_585_0_1 | *kiaa1429*-cluster #2 R | CCTCATTTCCCACGGTATCA |

### Supplementary Note 2. Measurement of m6A level in purified RNA

m6A RNA methylation level was measured by an m6A RNA methylation assay kit (Abcam, ab185912) following the manufacturer's protocol. Total of 200 ng of RNA isolated from control, *mettl14* (RNAi) and *kiaa1429* (RNAi) animals was used to determine the fraction of m6A in the RNA. Separately, m6A was measured on polyadenylated RNA and non-polyadenylated RNA from wild type planarians. polyA$^{+}$ RNA was separated from polyA- RNA using the NEXTFLEX Poly(A) Beads 2.0 kit (PerkinElmer). Following anti-m6A-antibody capture, the absorbance was measured at 450 nm using a microplate reader, and the percentage of m6A in the RNA was calculated using the kit supplied controls.

### Supplementary Note 3. RNA extraction

Samples were collected into TRI Reagent (Sigma; CAT #9424) and then homogenized using 0.5 mm zirconium beads in a bead beating homogenizer (Allsheng; Bioprep-24) for 2 cycles of 45 seconds at 3500 rpm followed by incubation of 5 minutes at room temperature. Then, 140 µl of chloroform was added to each tube. Tubes were shaken for 15 seconds, and then incubated at room temperature for 3 minutes. Then, samples were centrifuged at 4°C, 12000G for 25 minutes. The upper phase was separated into a new tube. Then, 500 µl of isopropanol was added, and tubes were inverted 5 times, and then incubated for 10 minutes at room temperature. Samples were centrifuged at 4°C (12000G) for 45 minutes. The supernatant was removed and the pellet was washed twice with 75% EtOH, centrifuged at 4°C 7500G for 5 minutes, and air-dried for 10 minutes. RNA was resuspended in 30 µl of $H_2O$. RNA concentration was measured by Qubit (Invitrogen; Q33226) according to the manufacturer’s protocol.

### Supplementary Note 4. mRNA m6A immunoprecipitation library preparation

m6A immunoprecipitation was performed as previously described[1]. RNA samples were extracted from *kiaa1429* (RNAi) animals or control, *unc22* (RNAi), animals in triplicates, following five dsRNA feedings. Each replicate included at least nine animals. RNA was quantified using Qubit (Invitrogen, Q33226) and 20 µg of *kiaa1429* (RNAi) and 30 µg of *unc22* (RNAi) RNA were used per sample per replicate. mRNA was enriched by 2 rounds of polyA selection (Dynabeads, CAT #61011) according to the manufacturer's protocol. mRNA-enriched samples were quantified by spectrophotometry (SpectraMax QuickDrop, Molecular Devices) and fragmented by RNA Fragmentation Reagents (Ambion, AM8740) for two minutes. Fragmented RNA was purified by MyOne Silane beads (Thermo, 37002D). Then, the purified RNA was treated with Turbo DNase (Thermo Fisher Scientific, AM2238) and the RNA was treated with FastAP (Fermentas, EF0651) and T4 PNK (CAT M0201L), according to the m6A pulldown protocol. Each RNA sample was barcoded by ligation to a unique 3’ adapter, which was flanked by a universal sequence that was used subsequently for reverse transcription. Then, the barcoded RNA samples were pooled by mixing and 10% of the pooled sample volume was separated and used as control (input) for the m6A pulldown experiment. Immunoprecipitation of m6A was performed by using an anti-m6A-antibody (Synaptic Systems) according to protocol. RNAseq libraries were prepared from the immunoprecipitated RNA and input RNA, separately, following the published protocol[1]. Library integrity was analyzed by fragment analysis (Agilent TapeStation 4000). Libraries were sequenced on an Illumina (NextSeq 550) machine at the Tel Aviv University NGS unit.

### Supplementary Note 5. Processing of m6A-seq2 library data

m6A-seq2 libraries were pooled and sequenced in paired-end mode[1]. The resultant sequencing reads were then demultiplexed (i.e., separated to individual libraries) based on an online library adapter that was ligated to each library according to the protocol[1]. Demultiplexing was performed by using cutadapt --pair-filter=any --minimum-length 24[2] by supplying internal barcode sequences, followed by trimming of the 10 nt sequence of the barcode from the barcoded sequencing read (Read #2). The resultant fastq files were pre-processed and mapped to the planarian transcriptome as described in the RNAseq library analysis method section. m6A-enriched regions (m6a peaks) were determined using MeTPeak with default parameters[3]. Plots of m6A peak profiles and peak heatmaps were produced using deepTools2[4]. Peak profiles were generated by merging BAM files of the three replicates of the control or *kiaa1429* (RNAi) libraries using the samtools merge command[5]. Then, mapping coverage was calculated for the merged control or *kiaa1429* (RNAi) BAM file using the deepTools2 command bamCoverage with parameters [--normalizeUsing RPKM --binSize 30]. The resultant files were then used for generating peak matrices with the computeMatrix scale-regions command, which was followed by producing the profile plots with the plotProfile command with parameters [--plotType lines --perGroup]. Peak heatmaps were produced by taking all of the detectable m6A peaks that were enriched by at least 5 folds in the pulldown libraries over the input. Then, the computeMatrix scale-regions command was used followed by the command plotHeatmap with parameters [--kmeans 3 --t "#2166ac,#d1e5f0,#ef8a62,#b2182b”].

#### Inline m6a-seq2 barcode sequences

| Condition | Replicate | Barcode sequence |
|---|---|---|
| *kiaa1429* (RNAi) | #1 | NNNCCAGTCT |
| *kiaa1429* (RNAi) | #2 | NNNCCTCCGT |
| *kiaa1429* (RNAi) | #3 | NNNGCACTCT |
| *unc22* (RNAi), Control | #1 | NNNGCCCATT |

| | | |
|---|---|---|
| *unc22* (RNAi), Control | #2 | NNNGGCCTCT |
| *unc22* (RNAi), Control | #3 | NNNTATCACT |

**Mapping of library data to the planarian transcriptome**

| Condition | Type | Mapping ratio | Unmapped read ratio |
|---|---|---|---|
| *kiaa1429* (RNAi) | Input | 0.871783 | 0.128217 |
| *kiaa1429* (RNAi) | pulldown | 0.876082 | 0.123918 |
| *kiaa1429* (RNAi) | Input | 0.872294 | 0.127706 |
| *kiaa1429* (RNAi) | pulldown | 0.88221 | 0.11779 |
| *kiaa1429* (RNAi) | Input | 0.88312 | 0.11688 |
| *kiaa1429* (RNAi) | pulldown | 0.885548 | 0.114452 |
| Control | Input | 0.886476 | 0.113524 |
| Control | pulldown | 0.895546 | 0.104454 |
| Control | Input | 0.840594 | 0.159406 |
| Control | pulldown | 0.891526 | 0.108474 |
| Control | Input | 0.882362 | 0.117638 |
| Control | pulldown | 0.893802 | 0.106198 |

### Supplementary Note 6. qPCR analysis of gene expression following RNAi

RNA concentration of each sample was measured using Qubit 4 fluorometer (Invitrogen; Q33226). cDNA synthesis was performed on 1 µg of RNA from each sample using RevertAid H Minus First Strand cDNA Synthesis Kit (Thermo scientific; K1631), following these steps: RNA template was mixed with an oligo-dT primer, and then incubated at 65°C for 5 min. Then, cDNA synthesis proceeded according to the manufacturer's protocol: samples were incubated at 42°C for 60 min, and then at 70°C for 5 min. The cDNA sample concentrations were normalized to 1 ng/µl. The expression of the target genes was measured using the QuantStudio 3 Real-Time PCR system (Applied Biosystems) using the following program [95°C (20 s), 40 cycles (95°C 1 s, 60°C 20 s)], with two technical replicates per sample and at least 2 biological replicates per sample. The relative gene expression fold-change was calculated by the ΔΔCt method with *gapdh* used as an endogenous control. Forward and reverse primers were designed for each target gene. Primer efficiency was tested prior to the experiment by using the standard curve method for 5 decreasing cDNA concentrations.

**qPCR primers used in this study**

| **Gene** | **Sequence** | **Contig** |
|---|---|---|
| *11930*_F | GAGGATTCTCCTATTAAACCCAAC | dd_smed_v6_11930_0_1 |
| *11930*_R | CGTGCGATCAGCGTTTATATT | dd_smed_v6_11930_0_1 |
| *gapdh*_F* | TCTTCCCCAACCAATTTTCTGTTCTG | |
| *gapdh*_R* | CCGAATATTTTATTTGGCTCTTCCTCCA | |
| *smedwi1*_F* | GTCTCAGAAAACAACTAAAGGTACAGCA | |
| *smedwi1*_R* | TGCTGCAATACACTCGGAGACA | |

| | | |
|---|---|---|
| *mettl3*_F | ACAGTTTTGTCACTATGGTACC | dd_Smed_v6_6450_0_1 |
| *mettl3*_R | GAAAGCAAGTATTGAGAAATGAAC | dd_Smed_v6_6450_0_1 |
| *wtap*_F | GGAGCTGATTTCCATTAAGAATG | dd_Smed_v6_2322_0_4 |
| *wtap*_R | TCGATATTGATTTTGCACACTG | dd_Smed_v6_2322_0_4 |
| *3194*_F | CATTTGTGCTATTACGGTGTC | dd_smed_v6_3194_0_1 |
| *3194*_R | GGTGAGTTCATTAGTGTACAGC | dd_smed_v6_3194_0_1 |
| *h2b*_F | TGTAAGGTTGATTTTACCAGGAG | dd_smed_v6_4808_0_1 |
| *h2b*_R | CCCGTGTACTTAGTAACAGC | dd_smed_v6_4808_0_1 |
| *rbm15*_F | GGCGAATTCAATGTCAAAGTAG | dd_Smed_v6_7282_0_1 |
| *rbm15*_R | AGCCAAAACAGGATCAACTG | dd_Smed_v6_7282_0_1 |

* Primer sequences previously published[6].

### Supplementary Note 7. Analysis of RNA sequencing gene expression changes

RNAseq sequencing files were preprocessed by trimming the Illumina adapter sequences using trimmomatic V0.38[7] [ILLUMINACLIP:TruSeq3-PE.fa:2:30:10:2:keepBothReads LEADING:0 TRAILING:3 SLIDINGWINDOW:4:15 MINLEN:30]. Then, the trimmed RNAseq sequences were mapped to the planarian transcriptome (ddV4)[8] and genome[9] using bowtie2[10] and hisat2[11], respectively. Gene expression matrix was then produced using the featureCounts command from the subread-2.0 package with parameters [-M -s 0][12] for the analyzed RNAseq libraries. Differentially expressed genes were detected using edgeR (3.30.3) with functions glmQLFit and glmQLFTest[13], and with DESeq2 (1.28.1)[14].

### Supplementary Note 8. Detection of polyadenylated transcripts in RNAseq data

Paired-end RNAseq libraries were preprocessed with trimmomatic V0.38[7] with parameters [ILLUMINACLIP:TruSeq3-PE.fa:2:30:10:2:keepBothReads LEADING:0 TRAILING:3 SLIDINGWINDOW:4:15 MINLEN:30]. For RNAseq libraries, reads in the Read #2 files were derived from antisense paired-reads. Sequencing reads from Read #2 files, which contained homopolymers (>10 nt) were discarded, but their read pair (Read #1) was kept. The retained Read #1 file was then mapped to the planarian transcriptome[9] using bowtie2 V2.3.5.1 with default parameters[10]. Reads that mapped to histone-encoding genes were isolated from the resultant read mapping BAM file[5]. Then, the unmapped read-pair (Read #2) of the isolated mapped-reads (Read #1) were selected from the raw paired-end RNAseq fastq files. An unmapped Read #2 sequence that included at least 10 consecutive T nucleotides (i.e., the reverse complement of the polyA) was counted as a polyA-containing read. The number of polyA-containing reads was summarized per library, and normalized by the library size.

### Supplementary Note 9. Quantification of polyadenylated *h2b* fraction by qPCR

RNA was isolated from *kiaa1429* (RNAi) and from control animals. Purified RNA was converted to cDNA by using random hexamers or poly dT primers. The cDNA samples concentrations were normalized to 1 ng/µl. qPCR was performed using QuantStudio 3 Real-Time PCR system (Applied Biosystems) with the following settings [95°C (20 s), 40 cycles (95°C 1 s, 60°C 20 s)], with two technical replicates per sample and at least 2 biological replicates per sample. The relative fold gene expression was calculated by the ΔΔCt method with *gapdh* used as the endogenous control.

### Supplementary Note 10. Preparation of scRNAseq libraries following *kiaa1429* (RNAi)

Animals were fed 5 times with *unc22* (control) and *kiaa1429* dsRNA. Animals were starved for a week following RNAi feedings. Cells were purified for scRNAseq from either control or *kiaa1429* (RNAi) animals by FACS using hoechst-labeling, as previously described[15]. Cells were counted and collected by centrifugation (300G, 5 min). The supernatant was discarded, and the cells were resuspended in 1x PBS with 0.5% BSA to achieve an optimal cell concentration of approximately

1000 cells per µl. Cells were counted automatically using the Invitrogen Countess II (Thermo Fisher Scientific) and the cell viability was assessed by labeling with trypan blue (0.4%) and counting positive cells.

Libraries were then prepared at the Genomic Research Unit at Tel Aviv University using the 10x Genomics Chromium Controller in conjunction with the single-cell 3′ v3.1 kit, protocol revision D. Briefly, cell suspensions were diluted in nuclease-free water according to the manufacturer's protocol to 10,000 cells in each sample. The cDNA synthesis, barcoding, and library preparation were carried out according to the manufacturer's instructions. cDNA libraries were sequenced on Illumina NextSeq 550 at Tel Aviv University.

### Supplementary Note 11. Systematic gene expression analysis of published planarian RNAseq data

A table of previously published planarian high-throughput sequencing libraries was extracted from the Short Read Archive[16]. The table (Supplementary Table 4) was manually curated and libraries were divided into biological groups, in each experiment, based on the library description in the Short Read Archive. Then, each library was downloaded using sra-tools[16] and was processed on a computational cluster using the computational pipeline used for the RNAseq libraries that were produced in this project. The resultant differential gene expression analysis tables, which were produced for the previously published planarian data, were used for searching for the genes that were differentially expressed in the libraries produced in this project with the following thresholds (FDR < 0.0001, $\log_2 > 1$ or $\log_2 < -1$).

### Supplementary Note 12. Analysis of human and mouse gene expression data from *CHD4* inhibition or conditional knockout

RNA-seq libraries of control and conditional knockout of *CHD4* from mouse[17], and from control and shRNA-treated samples against *CHD4* from human[18] were obtained from the short read archive using the fasterq-dump command[16]. The raw fastq files were trimmed using Trimmomatic-0.38[7] [ILLUMINACLIP:TruSeq3-PE.fa:2:30:10:2:keepBothReads LEADING:0

TRAILING:3 SLIDINGWINDOW:4:15 MINLEN:30]. Mouse and human samples were mapped to genome assemblies mm10 and hg38, respectively, using hisat2 with default parameters[11]. Gene expression was estimated using featureCounts[12] using the included human or mouse annotations. Finally, differential gene expression analysis was performed using DESeq2[14] with the DEseq pipeline. Resultant differential gene expression tables were annotated with the bioconductor annotation packages org.Mm.eg.db and org.Hs.eg.db, for mouse and human, respectively[19].

**Supplementary Table 1. m6A-enriched regions across the planarian transcriptome**

**Supplementary Table 2. Gene expression changes following inhibition of m6A genes**

**Supplementary Table 3. scRNAseq of *kiaa1429* (RNAi) and control animals**

**Supplementary Table 4. Published planarian high-throughput sequencing libraries compared to libraries produced here**

**Supplementary Table 5. Re-analysis of *CHD4* (RNAi) RNAseq from Tu et al., 2015**

**Supplementary Figure 1**

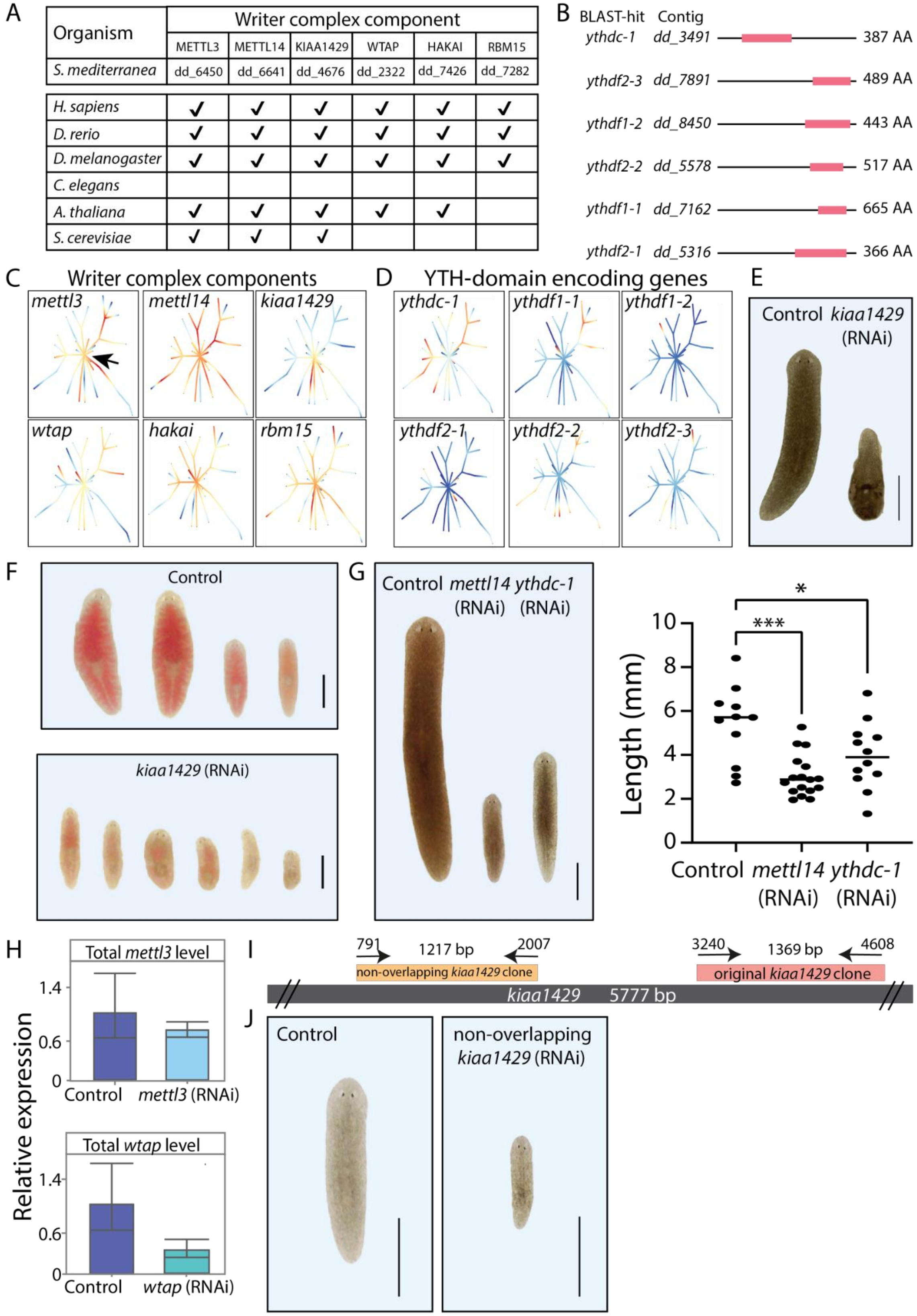

**Supplementary Figure 1. m6A genes are conserved and functional in planarians. (A) Conservation of the genes encoding the MTC is shown across diversity of organisms. Annotation for *A. thaliana* was obtained from a previous analysis[20]. (B) Domain analysis of putative YTH-domain (red block) containing genes. (C-D) The gene expression of planarian MTC and reader-encoding genes, across different cell types and lineages, was extracted from the planarian scRNAseq resource[21] (blue to red, low and high expression levels, respectively). The expression of MTC-encoding genes is widespread in different cell types, including in the stem cell compartment (black arrow, indicated in the *mettl3* panel). The expression of reader encoding genes is not widespread, except for *ythdc-1*, which is expressed across multiple cell types and conditions. (E) Inhibition of m6A genes resulted in lysis of the animals and eventually death. Shown are representative control and *kiaa1429* (RNAi) animals. (F) Inhibition of m6A genes resulted in defects in food uptake. Shown are animals following feeding with calf liver mixed with a red food color. Control animals show normal food uptake (top) compared with *kiaa1429* (RNAi) animals, which stopped eating. (G) Inhibition of *mettl14* and *ythdc-1* by RNAi has resulted in size reduction. Shown are representative images (left) and measurement of animal sizes following nine RNAi feedings. Asterisks represent p-value (One-way ANOVA followed by Dunnett's multiple comparison test; *** < 0.001; * < 0.05). (H) qPCR analysis has shown that the gene expression of *mettl3* (top) and *wtap* (bottom) is not downregulated significantly following RNAi, which could likely explain the lack of penetrant phenotypes in these conditions. (I) Schematic of the design of two non-overlapping gene fragments that were used for synthesis of dsRNA targeting the *kiaa1429* gene. The sequence used for the experiments presented in the manuscript was labeled "original *kiaa1429* clone". (J) dsRNA that was produced using a second cloned fragment of *kiaa1429*, labeled "non-overlapping *kiaa1429* (RNAi)", produced phenotypes that were similar to the phenotypes observed in a non-overlapping clone. This further indicated that the *kiaa1429* (RNAi) phenotype resulted from inhibition of *kiaa1429* gene expression and not because of an off-target effect of the RNAi.**

241 **Supplementary Figure 2**

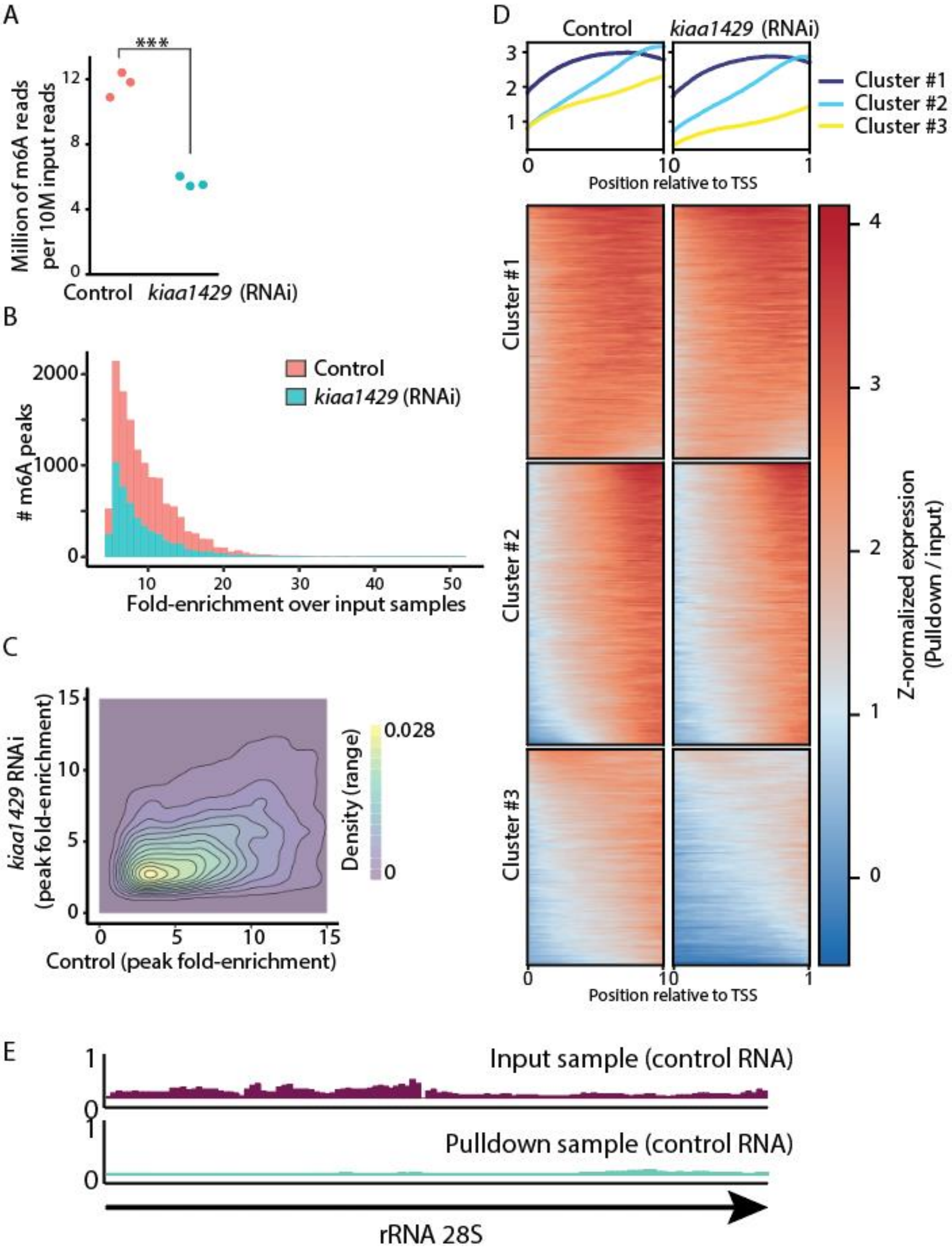

**Supplementary Figure 2. m6A is abundant on planarian mRNA and depleted following *kiaa1429* RNAi. (A) The number of RNAseq reads for each m6A-seq2 pulldown library per 10M of the corresponding input reads is shown. The number of reads in each pulldown library in the m6A-seq2 protocol[1] is correlated with the abundance of m6A (red and blue dot, control and *kiaa1429* (RNAi) sample, respectively. *** - Student's t-test $p < 0.001$). (B) The number of m6A-rich regions (peaks) is larger in control samples and depleted following inhibition of *kiaa1429*. Moreover, the fold-enrichment of the peaks in the control samples over the gene expression observed in the input samples is greater, in comparison to the *kiaa1429* (RNAi) samples. (C) A 2d-density plot showing the correlation between m6A-enriched regions in control and *kiaa1429* (RNAi) animals. The density plot shows that m6a peaks are more highly enriched in the control samples compared to *kiaa1429* (RNAi) samples, demonstrating the depletion of m6A following inhibition of *kiaa1429*. (D) Profile of m6A-enriched regions across genes in control and *kiaa1429* (RNAi) shows enrichment towards the 3'-end. The length of transcripts with detectable m6A-enriched regions was normalized to 1000 bp. Then, the expression across the transcript was computed by generating bins of gene expression (Supplementary Note 5) and calculating the log-fold change between the anti-m6a-antibody pulldown library and the input sample. K-means was used to separate three profiles of gene expression. (E) Shown is the RPKM normalized expression of the planarian rRNA 28S (block arrow) in the m6A pulldown library and in control. There is no evidence that planarian rRNA is methylated with m6A.**

### Supplementary Figure 3

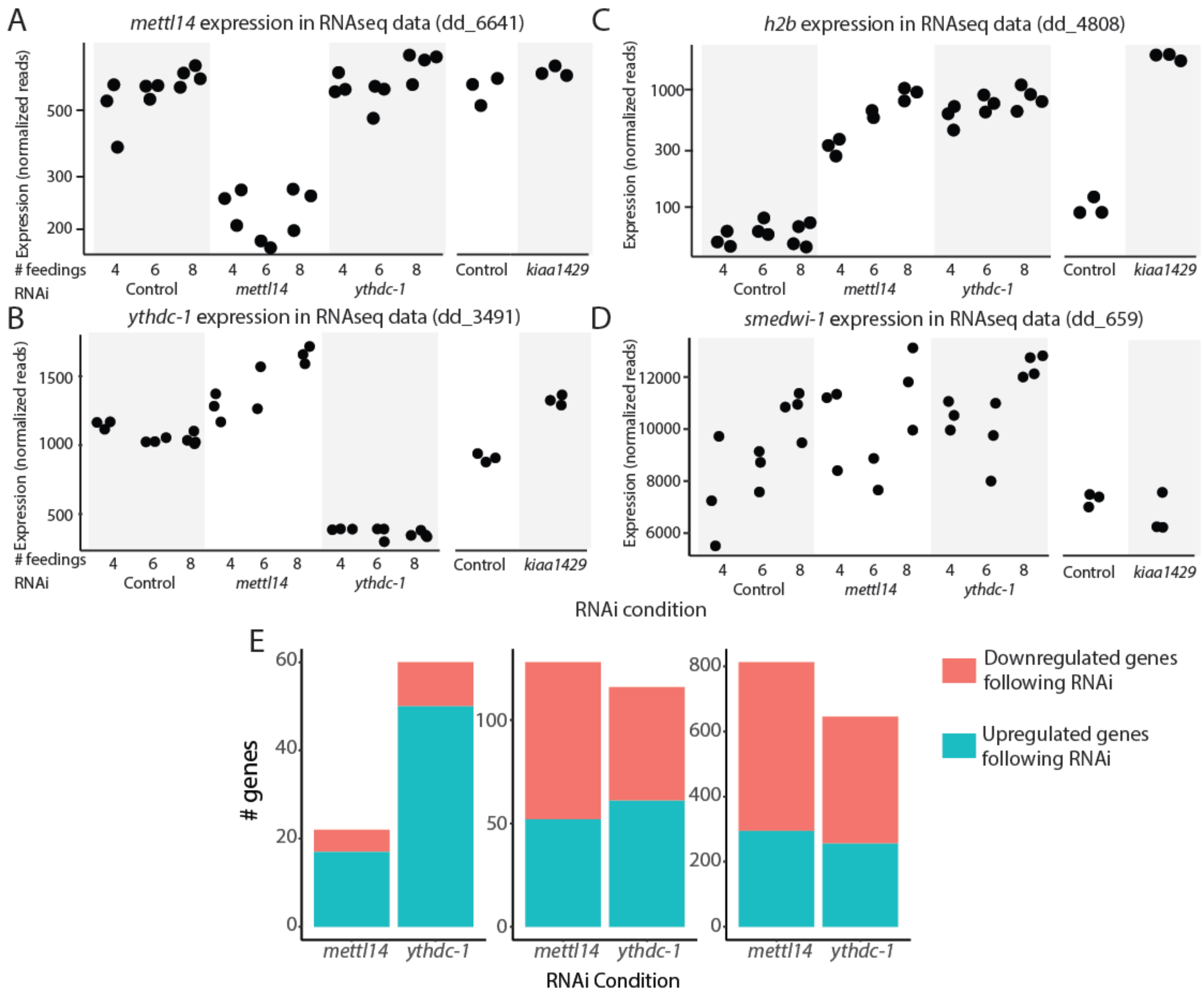


**Supplementary Figure 3. Specificity of RNAi of m6A genes. Shown is the normalized gene expression in RPKM for genes across the different libraries (A-D). RNAseq analysis validated the efficiency and the specificity of the RNAi (A-B). The expression of *h2b* was upregulated in all of the tested conditions (C). By contrast, the expression of *smedwi-1* did not change significantly following inhibition of m6A genes (D). (E) The number of significantly upregulated and downregulated genes is shown at each of the tested time points (FDR of genes included < 1E-5; Supplementary Table S2). In the early time point (four RNAi feedings) most of the genes were upregulated. However, at later time points, most differentially expressed genes are downregulated, which likely represent indirect effects of the RNAi, such as depletion of cell populations.**

### Supplementary Figure 4

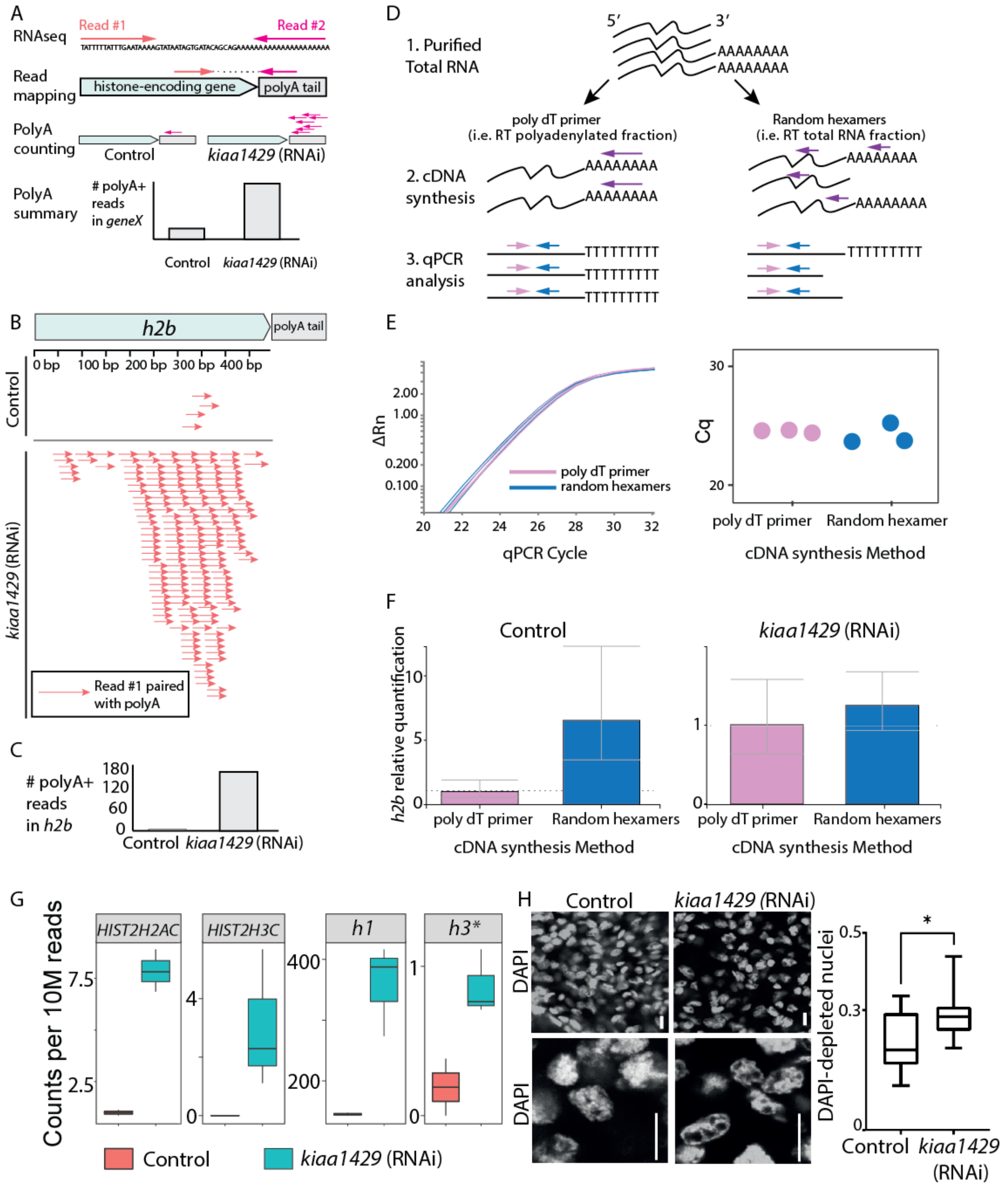


**Supplementary Figure 4. Analysis of histone transcript polyadenylation following inhibition of m6A genes. (A) Paired-end RNAseq data was used for identifying polyA containing histone transcripts. Reads mapped (read #1)**

**to histone genes without proper read-pairing were computationally isolated, and their read pair (read #2) was collected from the raw data files. If the unmapped read (read #2) contained a poly-dT sequence, which is the reverse-complement of polyA, then the read was counted as a polyadenylated histone transcript. (B-C) Shown are RNAseq reads mapped to *h2b* (dd_4808), which had a paired-read (read #2) containing polyA. Control sample (top) has fewer reads mapped to *h2b* compared to the *kiaa1429* (RNAi) sample (bottom). The total number of polyA containing read-pairs mapped to *h2b* is shown in panel C. (D) qPCR strategy for estimating the fraction of polyadenylated and non-polyadenylated *h2b*. Purified RNA (1) was reverse-transcribed with a poly-dT, or random hexamers (2). Then, qPCR is performed with internal primers (3). (E) The expression of *gapdh* was measured using RNA that was purified from wild type planarians. The analysis was used to estimate the efficiency of qPCR using cDNA produced either with poly-dT priming or using random hexamers. The change in reporter signal (y-axis) as a function of the qPCR cycle was similar for both groups (left), as is the quantification cycle (Cq, right). The analysis demonstrated that reverse-transcription with poly-dT and random hexamers was similarly efficient. (F) Shown in the relative quantification of polyadenylated *h2b* and non-polyadenylated *h2b* in control (left) and following *kiaa1429* (RNAi). The increase in polyadenylated *h2b* expression following *kiaa1429* (RNAi) was observed despite a lack of increase in total *h2b* expression (right). (G) Shown is the number of polyA containing reads for several of the overexpressed histone components encoding genes in RNAseq data per 10M library reads (*h3** is the putative *histone 3* transcript that is transcribed from contig dd_25629). (H) Comparison of DNA density by using DAPI-labeling on control and *kiaa1429* (RNAi) animals. DAPI-labeling, which correlates with chromatin density, showed an overabundance of DAPI-poor nuclei in *kiaa1429* (RNAi) animals. This suggested that *kiaa1429* (RNAi) chromatin packaging was altered, and indicated that euchromatin might be more prevalent in *kiaa1429* (RNAi), which may therefore affect transcriptional regulation. Scale = 10 µm.**

### Supplementary Figure 5

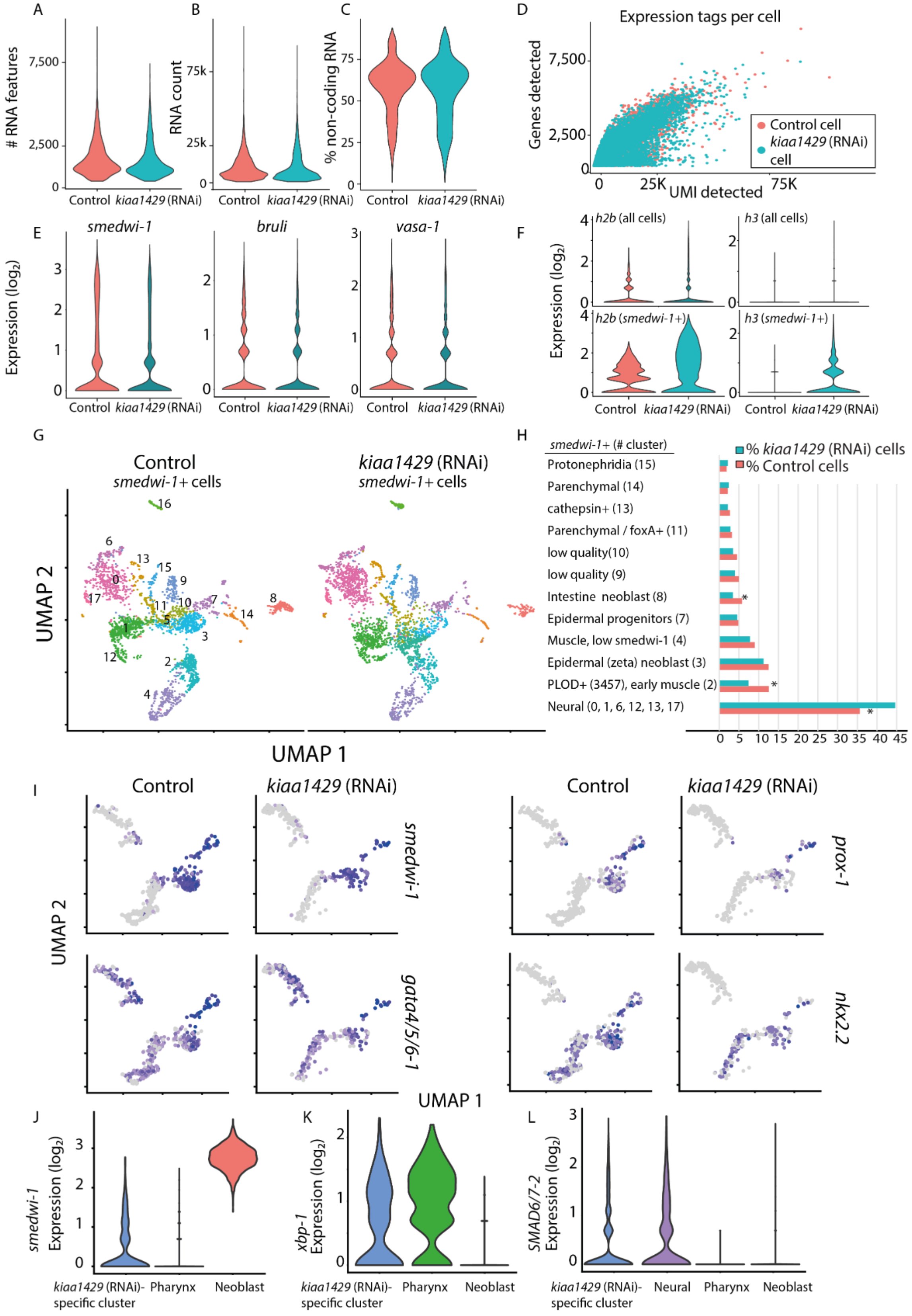

**Supplementary Figure 5. Analysis of *kiaa1429* (RNAi) inhibition by scRNAseq. (A-D) Quality measurements of scRNAseq libraries prepared from control and *kiaa1429* (RNAi) cells. The quality of the libraries produced from both conditions was assessed using the Seurat package[22]. The number of expressed genes (A); unique molecular tags (B); non-coding gene expression (C); and the correlation between the number of expressed genes and the unique molecular tags were highly similar between the libraries (D). This indicated that the quality of libraries was comparable. (E) Expression levels of canonical neoblast markers were highly similar in control and *kiaa1429* (RNAi). (F) The polyadenylated transcript expression of *h2b* and *h3* was much higher in *kiaa1429* (RNAi) animals compared to controls. The expression was detectable in neoblasts (bottom), and to a much lesser degree in the entire cell population (top), which includes neoblasts as well. (G-H) UMAP representation of neoblasts (color dots, left panel), and their identity (right panel). Neoblast identity was assigned based on expression of previously published gene expression markers[15]. Most neoblast clusters were not affected by the RNAi, yet several lineages (right, asterisk) were differentially represented following *kiaa1429* (RNAi). (I) Expression of neoblast (*smedwi-1*) and specialized intestine neoblast gene expression markers was overlaid on UMAP plots of the intestine lineage (cells represented by dots; grey and purple, low to high ranked expression). (J-L) Comparison of gene expression between several cell clusters. Importantly, pharynx cells are post-mitotic, and therefore show minimal *smedwi-1* expression (J). Post-mitotic cells express *xbp-1*, which was previously shown to be expressed in differentiating and differentiated cells[23] (K). *SMAD6/7-2* is expressed in the *kiaa1429* (RNAi) specific-cluster and in neural cells (L).**

### Supplementary Figure 6

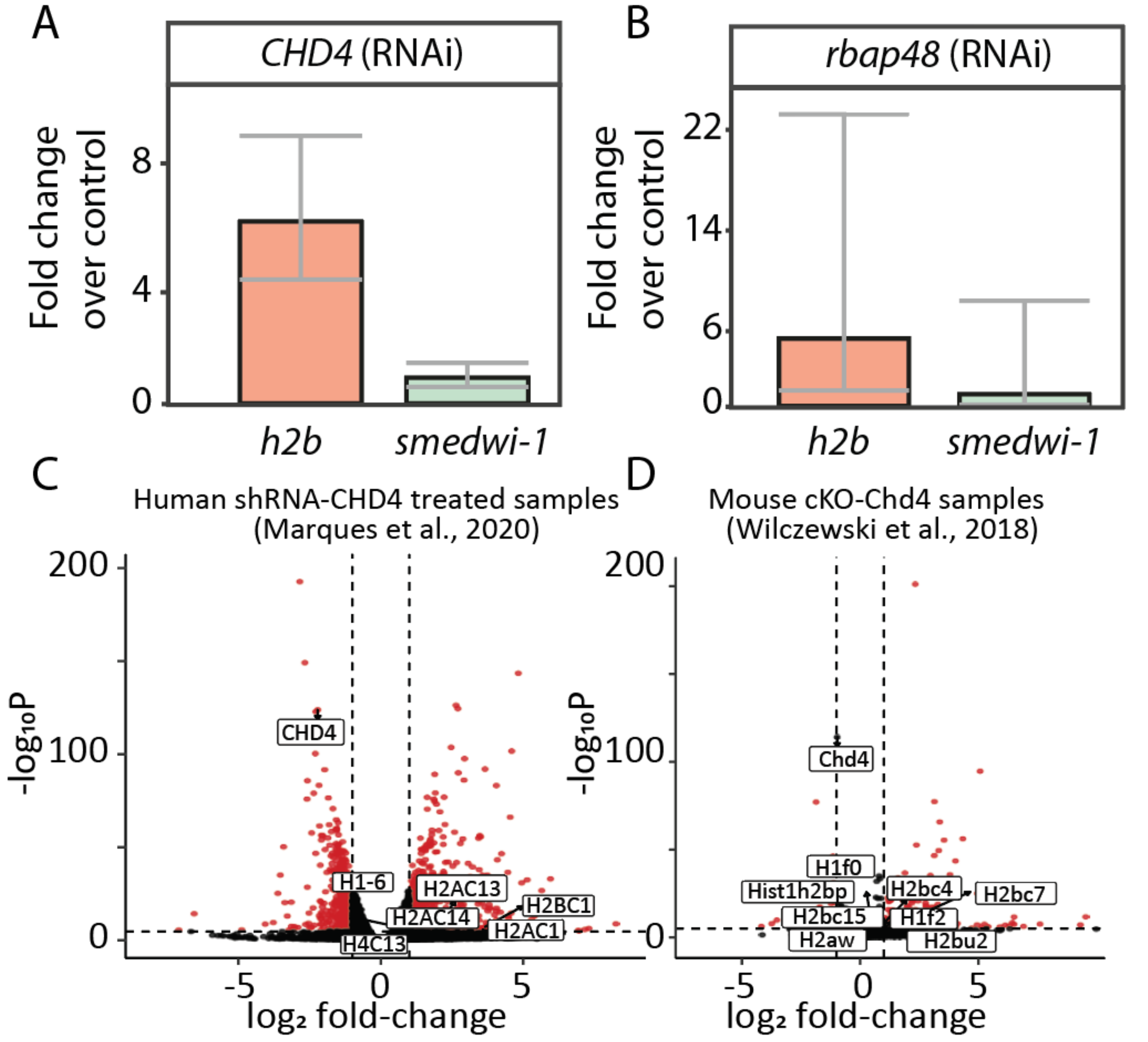


**Supplementary Figure 6. *CHD4* (RNAi) recapitulates molecularly the inhibition of m6A genes. (A-B) Shown is a qPCR quantification of *h2b* and *smedwi-1* expression following RNAi of the NuRD component encoding genes *CHD4* (A) or *rbap48* (B) in comparison to control libraries (error bars show 95% confidence interval). (C-D) volcano plot of human (C) and mouse (D) gene expression changes following inhibition or conditional knockout of CHD4 encoding gene. Highlighted and labeled are significantly overexpressed histone-encoding genes, as well as the human or mouse *CHD4* homolog (red and black, significant and non-significant change in gene expression; adjusted p-value < 0.0001; See Supplementary Note 12).**
